# Molecular dynamics of the pathogenic KCNQ2 variant G256W reveal mechanisms of channel dysfunction in epileptic encephalopathy

**DOI:** 10.64898/2025.12.01.691689

**Authors:** Emma C. Thompson, Edward C. Cooper, Bill R. Miller

## Abstract

Brain voltage-gated potassium channels containing the subunit KCNQ2 are essential for regulating electrical signals contributing to sensation, learning, memory, and motor control. De novo *KCNQ2* variants are among the more common Mendelian causes of early life epilepsy and neurodevelopmental impairment. Some patients with *KCNQ2* variants are affected with *KCNQ2* developmental and epileptic encephalopathy (*KCNQ2* DEE) characterized by seizures and developmental delays. Children with *KCNQ2* DEE exhibit a range of impairment patterns that appear to be correlated with specific consequences of the variant for protein function. Here, we used all-atom molecular dynamics to analyze a pathogenic missense variant KCNQ2 G256W, located in the pore turret. G256W subunit simulations showed migration of the hydrophobic W256 sidechain towards the lipid membrane. This movement affected turret structure and mobility prominently involving K255. We identified novel hydrogen bonding interactions in the wild type KCNQ2 turret region which formed a network that extended to the selectivity filter and identified N258, H260P, and K283 as key residues. Simulations comparing WT and G256W tetrameric channels exhibited more conformationally unstable ion selectivity filters for G256W. We analyzed how different stoichiometries of wild type and G256W subunits, as expected in heterozygous individuals, impacted dynamics and compared the G256W results to three additional variants of the turret-selectivity filter network. Our results provide additional support for an integral role of the KCNQ2 turret selectivity filter stability. The majority of severe *KCNQ2* DEE variants are clustered near the selectivity filter in the pore domain. Our study provides insights that may be broadly applicable to this clinically important allele subgroup.

**Broader audience statement:** A serious childhood illness, called *KCNQ2* developmental and epileptic encephalopathy, usually arises from single amino acid substitutions, or missense variants. This paper provides insight into how such a local change can profoundly disrupt the function of a large oligomeric protein containing over 3,400 residues. Molecular dynamics simulations of the pathogenic KCNQ2 pore domain variant, G256W, revealed that W256 changed the structure and flexibility of the pore domain turret and altered the ion selectivity filter. These findings shed light on the functional impact of KCNQ2 pathogenic variants and may help illuminate general mechanisms underlying severe KCNQ2 variants that occur commonly near the ion pore.

## 1. INTRODUCTION

Genetic variants of *KCNQ2* are among the most frequent Mendelian causes of epilepsy in children.^1,2^ There are many variants with different functional effects leading to a broad spectrum of *KCNQ2* epileptic and neurodevelopmental disorders. Milder variant effects cause self-limited familial neonatal epilepsy (SLFNE) characterized by pre and postnatal seizures up to a few months of life before stopping with neurotypical development.^3,4^ The more severe end describes profound developmental delays and treatment-resistant seizures called *KCNQ2* developmental and epileptic encephalopathy (*KCNQ2* DEE). This study focused on a particular variant in the middle of the spectrum, G256W, which presents with moderate developmental delays and neonatal onset epilepsy.^5^

The protein product of the *KCNQ2* gene called KCNQ2 or alternatively, Kv7.2, forms channels that contribute to the M-current, a low voltage-activated, non-inactivating, neuronal potassium current modulated by Gq-linked neurotransmitter receptors.^6–8^ G256W homozygous mice die at birth; G256W heterozygous mice show hyperexcitability in hippocampal pyramidal neurons, spontaneous convulsive seizures, premature death, and impaired protein localization to axon initial segment.^5,9^

Like other related voltage gated K^+^ (Kv) channels, KCNQ2 channels include transmembrane and large intracellular domains.^10,11^ Each subunit includes a voltage sensing domain (VSD) with four transmembrane helices (S1-S4), and a pore gating domain (PGD) with two-transmembrane segments (S5-S6) linked by the pore loop.^12–15^ The intracellular c-terminus is large and multimodular, and includes sites for binding calmodulin (CaM).^16^ The pore loop consists of three segments: the turret, pore helix and the selectivity filter. G256W is located near the S5 helix, at the beginning of the turret segment, a region known to be important for toxin-binding.^5,17^ After the selectivity filter, a segment (called the selectivity filter bridge) connects the transmembrane ion path to S6.^5^ The G256 α-carbon is about 22 Å from the extracellular opening of the selectivity filter.^5^ Why might substitution at this site so distant from the ion pore lead to loss of channel currents? Cryo-electron microscopy density maps provide evidence for a network of hydrogen bonds extending from G256 to the selectivity filter bridge.^14^ In this study, we use molecular dynamics simulations to test the idea that loss-of-function observed in G256W-containing channels arises from turret configurational changes that extend to the selectivity filter.

Computational modeling methods are often used to investigate biophysical effects of protein variants and explore structure-function relationships.^18^ Computational studies and new cryoEM structural datasets have recently been employed to analyze the effects of KCNQ2 channel activators such as retigabine and phosphatidyl inositol-4,5 bisphosphate (PIP2) that promote conduction through stabilization of the open state.^19–30^ Molecular dynamics (MD) simulations and analysis of KCNQ2 variants have been used to explain pathogenic consequences at the structural level.^21^

Here, using all-atom molecular dynamics, we measured protein stability and residue fluctuations, quantified non-covalent interactions in the pore domain, and compared solvent accessibilities and preferred configurations of G256 and W256 to understand the structural and functional consequences resulting from the *KCNQ2* pathogenic variant, G256W.

## 2. RESULTS & DISCUSSION

### 2.1 Turret and selectivity filter dynamics differ between WT and G256W

Native KCNQ2 channels are homotetrameric or heterotetrameric (with KCNQ3 and or KCNQ5).^7,31^ Here we mainly focus on homomers, channels containing four KCNQ2 subunits, although we briefly explore the differing WT:G256W stoichiometries later. Each polytopic subunit includes a voltage sensing domain, a portion of the channel’s pore domain, and large cytosolic domain; the extracellular domain is small, consisting of short loops (Figure 1A; Figure S1). G256 is located in the turret of the pore domain and was predicted by the cryoEM model to participate in a hydrogen bonding network.^5,27^ Figure 1B (WT) and 1C (G256W) indicate the location of substitution and illustrate the structural differences between the small glycine and planar, hydrophobic tryptophan.^32^ Notably, the extracellular ends of S5 and S6 both faced lipid and placed G256W far from voltage sensing domains.^33^ First, we performed simulations comparing wild type KCNQ2 tetramers to channels containing four G256W subunits. Although such channels represent a small fraction of the channels predicted to assemble within human patients who are heterozygous for the variant, this approach allowed us to collect data on four subunits per simulation.^5^

**Figure 1.**
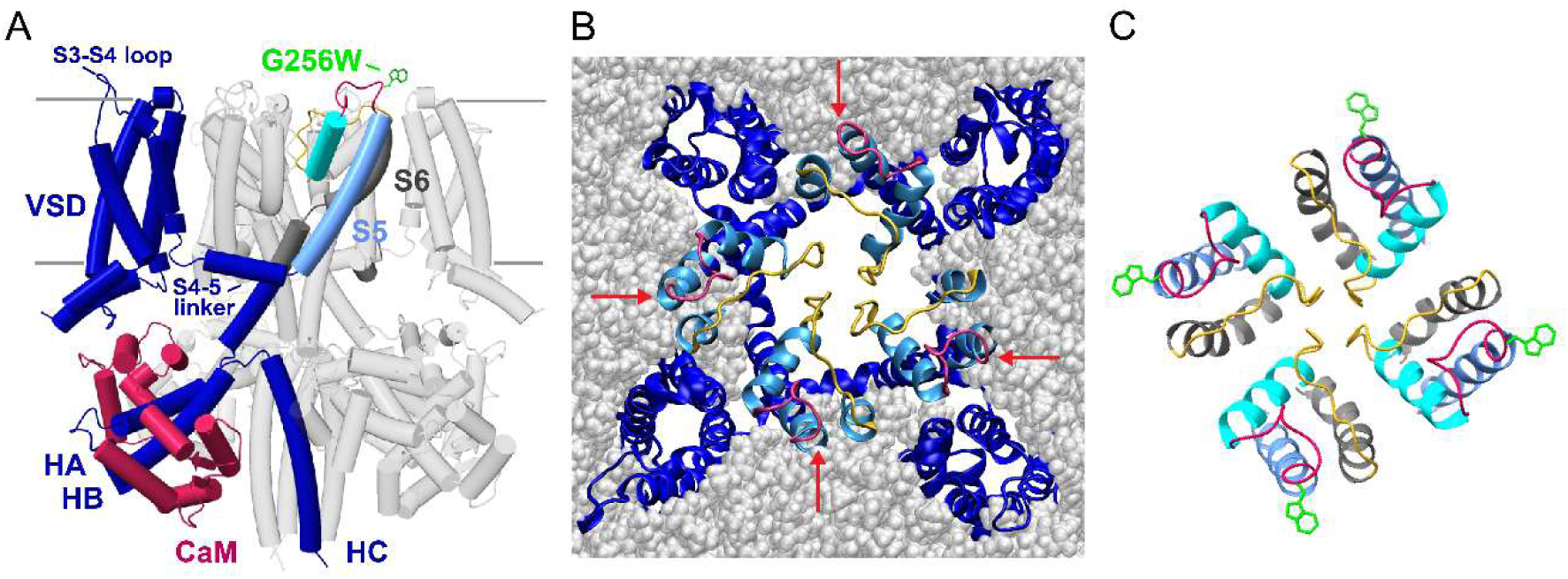
G256W lies in the extracellular turret region of the pore domain near the protein-lipid interface. A. Extracellular up, side view of a KCNQ2 tetramer based on the cryoEM model (PDB: 7CR3). One subunit, labeled in color, shows the modifications added in this study, i.e., the extracellular loop between S3-S4, and, near the top of the channel’s extracellular domain, the G256W substitution. Two subunits are in transparent gray; one subunit has been partially removed for clarity. **B.** Extracellular view of an MD simulation representative frame depicting a tetramer embedded in the membrane. Intracellular N and C terminal regions were omitted for clarity. Red arrows point to G256 locations, highlighting their proximity to lipid. **C.** Extracellular view of the tetrameric pore domain (PDB: 7CR3) extracellular region and transmembrane regions near the selectivity filter. The four substituted W256 sidechains are shown in green. Helix and segment colors are as shown for the colored subunit in **A**. Abbreviations: VSD, voltage-sensor domain; HA, HB, and HC, the cytoplasmic helices HA, HB, and HC; CaM, calmodulin.

Backbone atom root mean squared deviation (RMSD) of the triplicate simulation trajectories increased in deviation until, on average, stabilization near 250 ns (Figure S2). The standard deviation of RMSD for the 0 to 250 ns compared to 250 to 750 ns trajectory was substantially higher. To reach a stable low energy structure to model the structures likely to be sampled *in vitro*, all subsequent analysis focused on this post stabilization period 250 ns to 500 ns, termed the production run (Figure 2; Table S1). The WT production simulations average ± standard deviation (SD) RMSD was 5.98 ± 0.38 Å. The G256W protein RMSD average ± SD was 6.25 ± 0.52 Å. The Hedges’ g magnitude statistical difference between the three replicates for WT and G256W RMSD was 0.53, meaning there was a medium difference between them. With only four residues of difference between the two simulations, a medium effect size suggested that the simulations effectively differed in local structure. Individual averages and standard deviations for each simulation run (four subunits per run) are indicated in Figure 2. The G256W protein deviated more from the original structure, which was expected because all simulation runs began at coordinates from a WT cryoEM (PDB: 7CR3) model.

**Figure 2.**
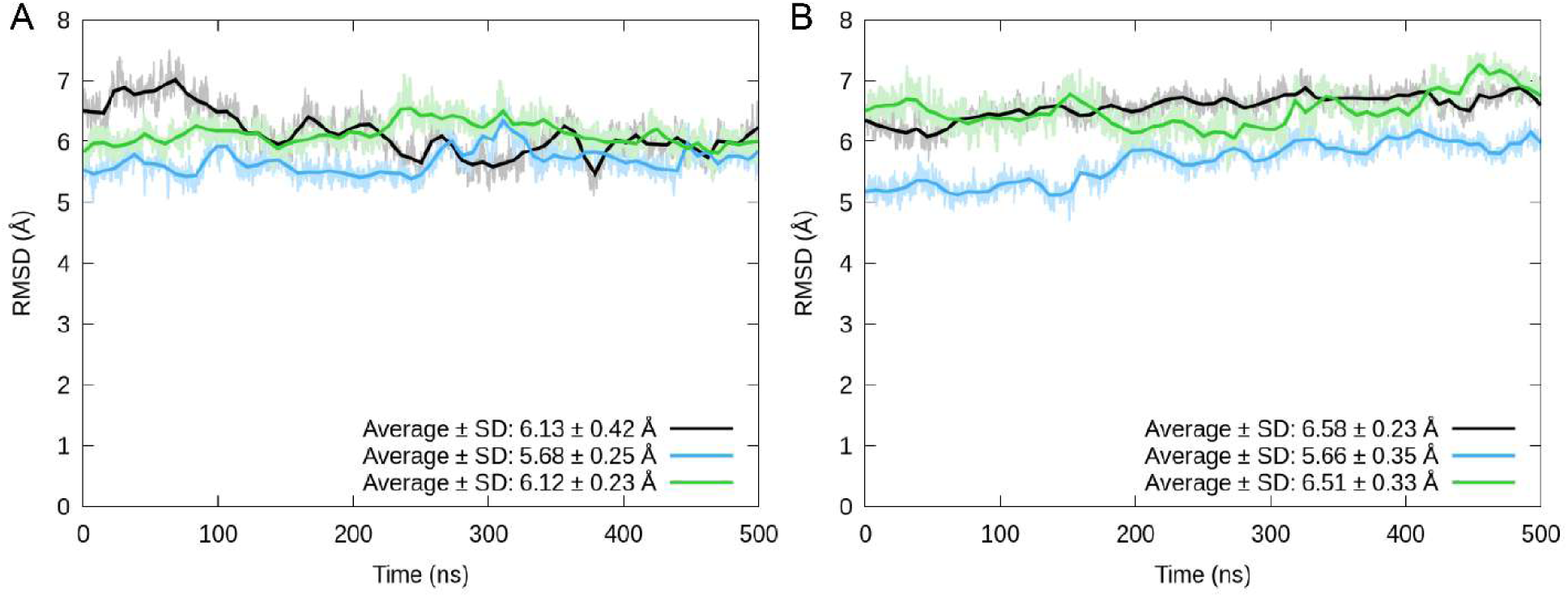
Root mean square deviation of the WT and G256W production simulations show structure stable structures and greater deviation for G256W. Root mean square deviation averaged across all subunits for **A.** WT and **B.** G256W 500 ns production simulations. Lines for replicates one (black), two (blue), and three (green) illustrate full structure RMSD traces (faint) and smoothed average lines (bolded) per simulation. Legend displays the calculated RMSD average and standard deviation for each replicate simulation.

The dynamics of the turret segment (residues 255 to 262) and the canonical selectivity filter (TI/VGYG, residues 277 to 281) were quantified using backbone RMSD (Figure 1; Figure S3). On average, the WT turret had access to more conformations across simulations than the G256W turret (Hedges’ g magnitude effect size between WT and G256W RMSD of 0.35). The average ± SD of the WT turret was 3.38 ± 1.02 Å. There was great variability amongst the WT replicates displayed. One of the WT replicates had the lowest average RMSD (2.05 Å) of all six simulations (three WT, three G256W) while two others had the highest turret RMSD (4.14 Å and 3.68 Å). Compared to G256W (average ± SD: 2.99 ± 0.44 Å), the WT turret on average had increased dynamics relative to the starting structure and a higher standard deviation which suggested more flexibility. A first comment is that these differences result from glycine, which does not have a side chain and therefore the backbone moves more freely.^34^ The selectivity filter followed the same trend as the turret; the WT (average: 3.01 Å) selectivity filter had a higher average RMSD than G256W (average: 2.46 Å). However, the G256W simulations showed a higher standard deviation (SD: 0.59 Å) for the selectivity filter than the WT (SD: 0.18 Å), which indicated that the variant had effects on ion pore dynamics (Hedges’ g magnitude effect size between WT and G256W of 0.8).

Residue root mean square fluctuation (RMSF) was calculated for only the polypeptide backbone atoms (“backbone RMSF”) and for all sidechain and backbone atoms (“sidechain RMSF”). Figure S4 compares backbone and sidechain RMSF observed in WT and G256W simulations. As expected, the sidechain RMSF has a higher fluctuation.

Backbone RMSF indicated that the WT and G256W overall protein structures were similar (Figure 3A). The backbone structure of the turret between the WT and G256W protein was similar as well (root mean square error, RMSE, of 0.043 Å; Figure S5; Table S2). Higher mobility for the solvent accessible S3-S4 loop (184-193) matched the flexibility of the homologous KCNQ1 region as described recently.^35^ The S3-S4 loop backbone RMSE (0.53 Å) between WT and G256W was 0.11 Å above the RMSE of the overall protein.

**Figure 3.**
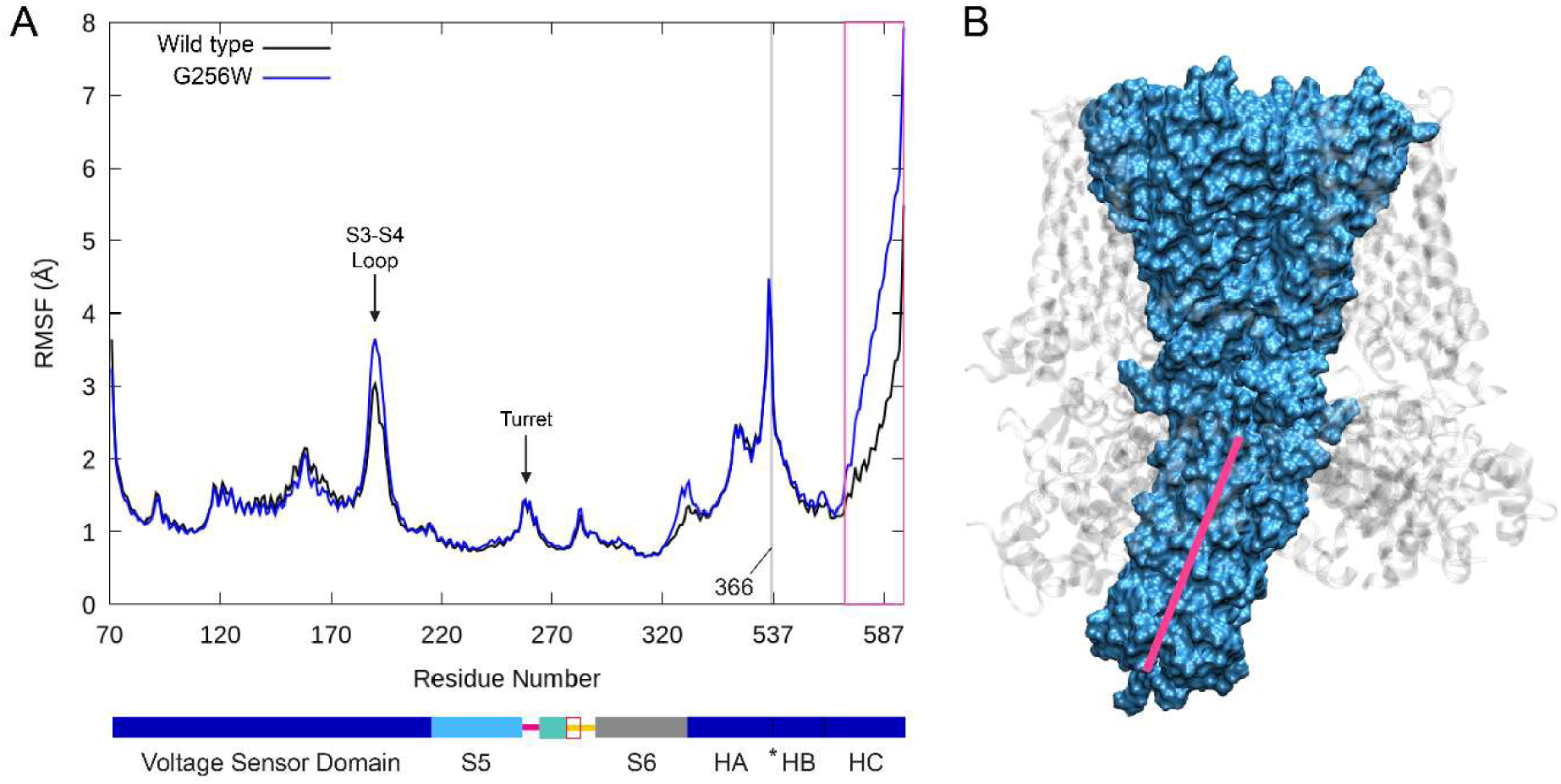
Preferential interaction between HC helix and one calmodulin subunit differing between WT and G256W indicates G256W may have long range structural effects. A. Backbone atom root mean square fluctuation averaged over all of WT (black) and G256W (blue) subunits. Arrows point to local peaks at S3-S4 loop and turret regions. A KCNQ2 domain map is shown below (map colors as in Figure 1A). The vertical grey line at residue 366 (asterisk in the domain diagram) indicates a jump in sequence to residue 536 corresponding to a poorly resolved HA-HB loop. **B.** Surface representation of the channel pore domain and HC helix. The more peripheral VSD and HA-HB-CaM components of KCNQ2 are transparent gray ribbons. A pink line highlights the asymmetric orientation of the tetrameric HC coiled-coil (corresponding to the pink box enclosed region in **A**).

The intracellular HC helix displayed a uniquely high backbone RMSF (Figure 3A, region boxed in pink). Examination of the tetramer structure associated with this revealed asymmetric deviation of the HC helix coiled-coil domain which led to interactions with calmodulin. A surface representation of the asymmetry (with a line that indicates the HC helix coiled-coil main axis) is shown in Figure 3B. The difference in fluctuation between WT and G256W across the length of HC revealed preferential interaction between the HC helix coiled-coil tetramer and one calmodulin subunit in G256W channels (RMSE 1.47 Å). It is important to note that these differences may be inflated due to more flexible artificial free termini because of missing HA-HB loop residues near the HC helix. Interestingly, however, such deviations from radial symmetry were previously observed using NMR solvation of the intracellular c-terminal fragments.^36^ This asymmetry was proposed to be a flexible conformational hinge that may affect channel gating properties.^36^

Side chain RMSF calculations provided evidence for distinct conformational dynamics between WT and G256W turrets (Figure 4). Whole sequence plot of the side chain RMSF revealed differences both near the variant and far from it, on both sides of the HA-HB helix gap in sequence of the intracellular domain (Figure 4A). The variability near the HA-HB helix gap is likely due to the presence of artificial free termini that are more flexible in the intact subunit, however it is not clear why this would differ between WT and G256W as we observed. Closer examination of the WT and G256W side chain RMSF plots near residue 256 revealed higher flexibility for the WT than G256W (Figure 4B). The turret RMSF indicated a higher flexibility for the WT side chains than G256W (RMSE: 1.62 Å), although similar flexibility in the backbone atoms (RMSE: 0.043 Å) revealed that the variant may affect conformational changes within turret side chains. Hedges’ g magnitude calculations between WT and G256W indicate a greater difference between G256W and WT turret RMSF including backbone and side chains than only backbone atoms for all residues, except D262 (Table S2). The largest difference with side chain RMSF was 3.2 Å at K255 (WT: 7.0 Å, G256W: 3.8 Å). In contrast, the backbone fluctuation of K255 was similar between the two structures (WT: 1.15 Å, G256W: 1.14 Å). This is consistent with the large difference between Hedges’ g magnitude differences between G256W and WT tetramers described in Table S2. The positively-charged K255 side chain may have fewer steric clashes when adjoining G256 than W256, which would explain its larger fluctuation in the WT turret. This was explored in more detail later.

**Figure 4.**
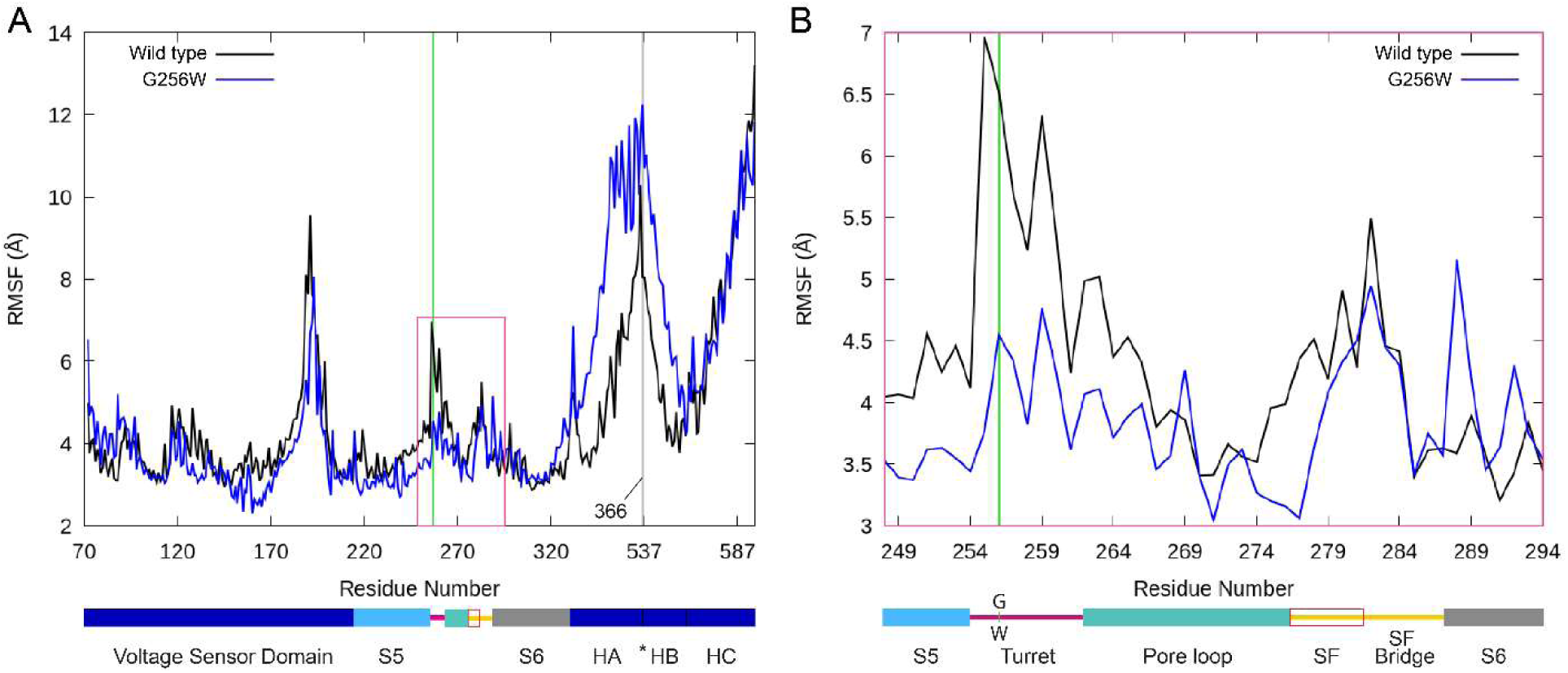
Root mean square fluctuation analysis reveals G256W shows reduced turret flexibility compared to the WT. Per residue root mean square fluctuation averaged over all subunits including side chains. Schematic representations of the KCNQ2 domains are shown below each graph with colors corresponding to Figure 1A. **A.** Comparison of the WT (black) and G256W (blue) structures of all modeled KCNQ2 residues. The grey line marks the point of omission of residues 367 to 535 (i.e., the HA-HB loop, unresolved in cryoEM structures). The green line marks G256W. **B.** A zoom-in on the pore domain region boxed in pink in panel **A**.

### 2.2 W256 side chain migration toward the membrane outer leaflet

Although simulation equilibration began with the G256 and W256 residues exposed to bulk solvent, calculation of solvent accessible surface area (SASA) percentage during the production simulations revealed a significantly reduced solvent accessibility of W256 compared to G256 (WT: 60.5% ± 0.2%, W256: 36.8% ± 0.1%; Welch’s t-test: t-statistic = 206.9, p-value < 0.001).^37^ The location of G256W at the periphery of the pore domain and at a distance from voltage sensor domains allowed for potential direct interactions with the outer leaflet of the lipid membrane (Figure 1B-C).Visualization of a sample of sidechain positions during the simulations revealed that burial of the large hydrophobic W256 side chain near the lipid-water interface accounts for the reduced calculated SASA. We extracted distances from the membrane center for subsets of atoms of G256 and W256 (Fig. 5C-E).^38,39^ The average distance from membrane center for the G256 and W256 backbone atoms are not different; the driver of W256 migration towards the membrane center is the side chain (Figure 5C-E). We compared subsets of side chain atoms to learn about the orientation of the side chain. The amine side of the planar indole has the smallest average distance (purple line, Figure 5C); however, all side chain atom subsets tested were significantly closer to the membrane center than the backbone. Variation of the membrane center distances of side chain atoms subset was highly coherent (Figure 5C). This membrane interaction by the W256 side chain may anchor the G256W turret, providing an explanation for observed less mobile turret compared to the WT (Figure 4B).

**Figure 5.**
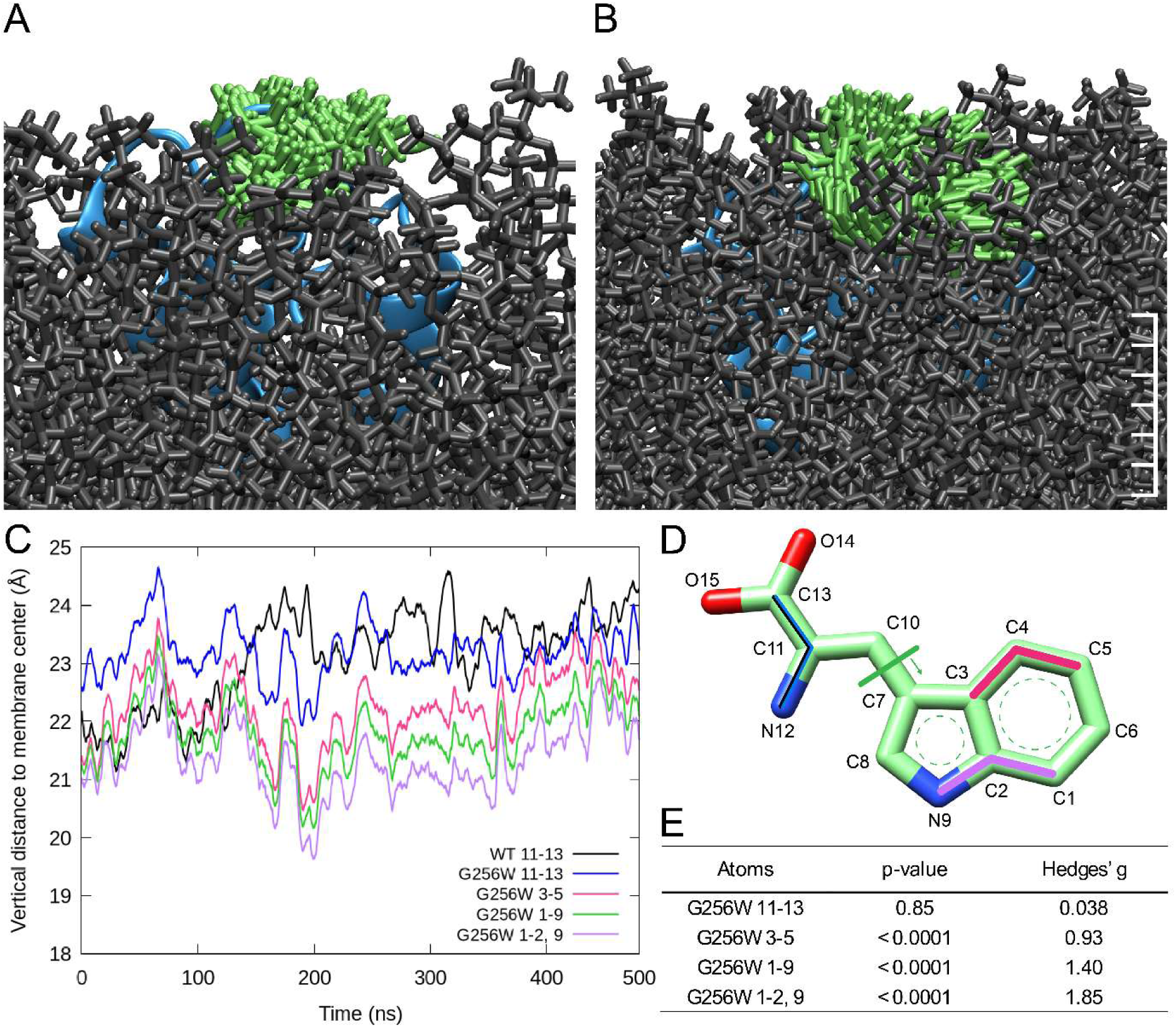
W256 side chain migration and burial into the lipid membrane. Superimposition of 150 images of single subunits from G256 and W256 simulations. **A.** G256 and **B.** W256 are colored green; samples are taken every 10 ns intervals over 1.5 µs. WT G256 is largely at or above the membrane surface, but W256 sidechain is often buried in the lipid (grey). Scale with tick marks are approximately 1 Å. **C.** Running average plots of the distance between membrane center and the average positions of G256 (black) or W256 (blue) backbone atoms, or selected subsets of W256 sidechain atoms. **D.** Trp stick model. Atom numbering corresponds to line plots in **C** and published literature.^38,39^ Distance to membrane center Average ± SD were 23.1 ± 1.1 Å (WT 11-13), 23.1 ± 0.9 Å (G256W 11-13), 22.2 ± 0.9 Å (G256W 3-5), 21.7 ± 0.9 Å (G256W 1-9), and 21.3 ± 0.9 Å (G256W 1-2, 9). **E.** W256 side chain atom subset distances are significantly smaller than the G256 mainchain distance.

### 2.3 Wild type turret flexibility allows for more available conformations and different interaction frequencies than G256W

Direct hydrogen bond frequencies of the WT and G256W simulations were calculated (Figure 6; Figure 7). Hydrogen bonds between N258 and E254, N258 and H260, and K283 with H260 predicted in Abreo et al’s interpretation of prior cryoEM data were each supported by the MD simulations, showing frequencies larger than 25%.^5,14^ In other respects, the MD network results were unlike the static cryoEM model; MD indicated that K255 was highly dynamic. The cryoEM density map distance predicted interaction between K255 and Y251 sidechains, but the simulations showed K255 interaction with Y251 and with 4 other turret residues (Figure 7A).

**Figure 6.**
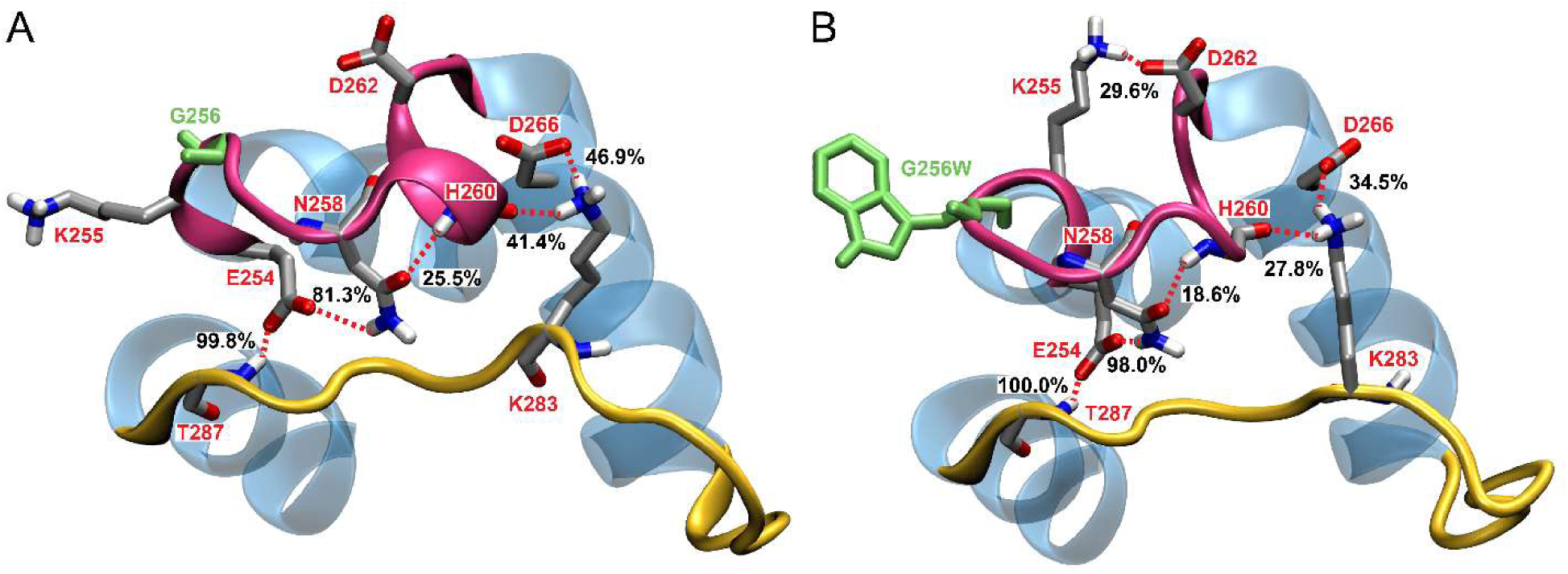
WT and G256W simulations reveal differences in turret network hydrogen bonding. Representative structure frames of **A.** WT and **B.** G256W turret regions are labeled for percentage of time spent hydrogen bonded to nearby residues across all simulations. G256 and W256 are shown in solid green; other amino acid sidechains are colored by heteroatom. S5, S6, and pore helices are shown in blue; the turret is pink, and the selectivity filter (including SF bridge) is shown in yellow. Prominent interactions and those captured in the selected still images are labeled.

**Figure 7.**
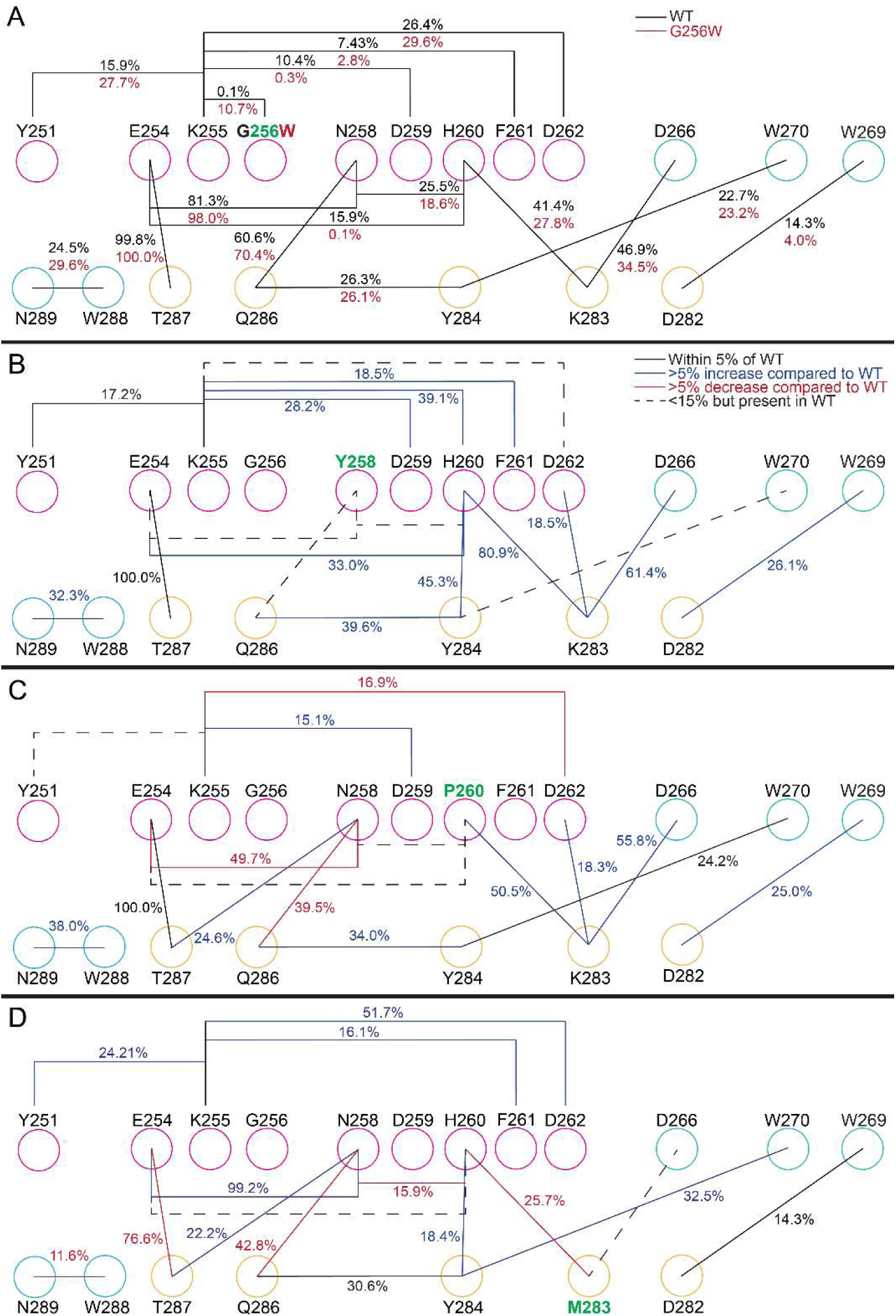
Network diagram of hydrogen bonding frequencies depict differences in WT and variant simulations. Hydrogen bonding network diagrams for the **A.** WT (black) and G256W (red) 1.5 µs of equilibrated simulations, and **B.** N258Y, **C.** H260P, and **D.** K283M. For **B-D**, percentages are averaged from the 200 ns simulations. Open circles are colored as pink (turret), teal (pore helix), yellow (selectivity filter bridge), and blue (helix S6). Labels on lines connecting residues display the percentage of frames showing interaction. The line color key in **B** (upper right) applies to **C** and **D**. Black lines represent hydrogen bonds within 5% of the WT simulations, blue and red lines indicate at least a 5% increase or decrease, respectively, compared to the WT. Dotted lines represent interactions in the variant simulations that are not present (less than 15%) in the WT.

Direct hydrogen bonds between backbone or side chain atoms of G256 and N258, H260 and Q286, and H260 and D262 predicted by the cryoEM model were not found in large percentages in the MD simulation. Importantly, both the cryoEM model and our WT simulations predicted three hydrogen bonds between the turret and selectivity filter bridge. In the simulations, the interactions were between residues N258 and Q286 (60.6%), E254 and T287 (99.8%), and D266 and K283 (46.9%) (Figure 6; Figure 7A), but only one of these (between D266 and K283) was seen in both the cryoEM model and MD simulations. These differences highlight additional residues and residue pairs for future experimental mutagenesis and electrophysiological characterization.

Analysis of the position and interactions of K255 revealed many differences between the WT and G256W simulations. Figure 8 shows top down and side views of K255 side chain positions sampled every 10 ns for WT and G256W simulations. WT K255 had access to a larger spread of conformational states with one subunit displaying a near bimodal overall distribution. The terminal amine of the WT K255 was brought proximate to the extracellular side of the pore helix where it may have formed a salt bridge with the carboxylic acid of D262. K255 also shifted closer to the S6 helix in the WT simulation, which was not seen in the G256W simulations (Figure S8). The W256 (sidechain in green) effectively blocked K255 from occupying conformations on the other side of the turret. A higher frequency of interaction with D259 in the WT structure was explained by the greater flexibility of K255’s side chain (Figure 7A). In the WT structure, K255 showed lower frequencies of interactions with D262, Y251, and G256 (Figure 6; Figure 7A). K255’s lower probability of these interactions was likely due to the WT having more conformations accessible (Figure S8).

**Figure 8.**
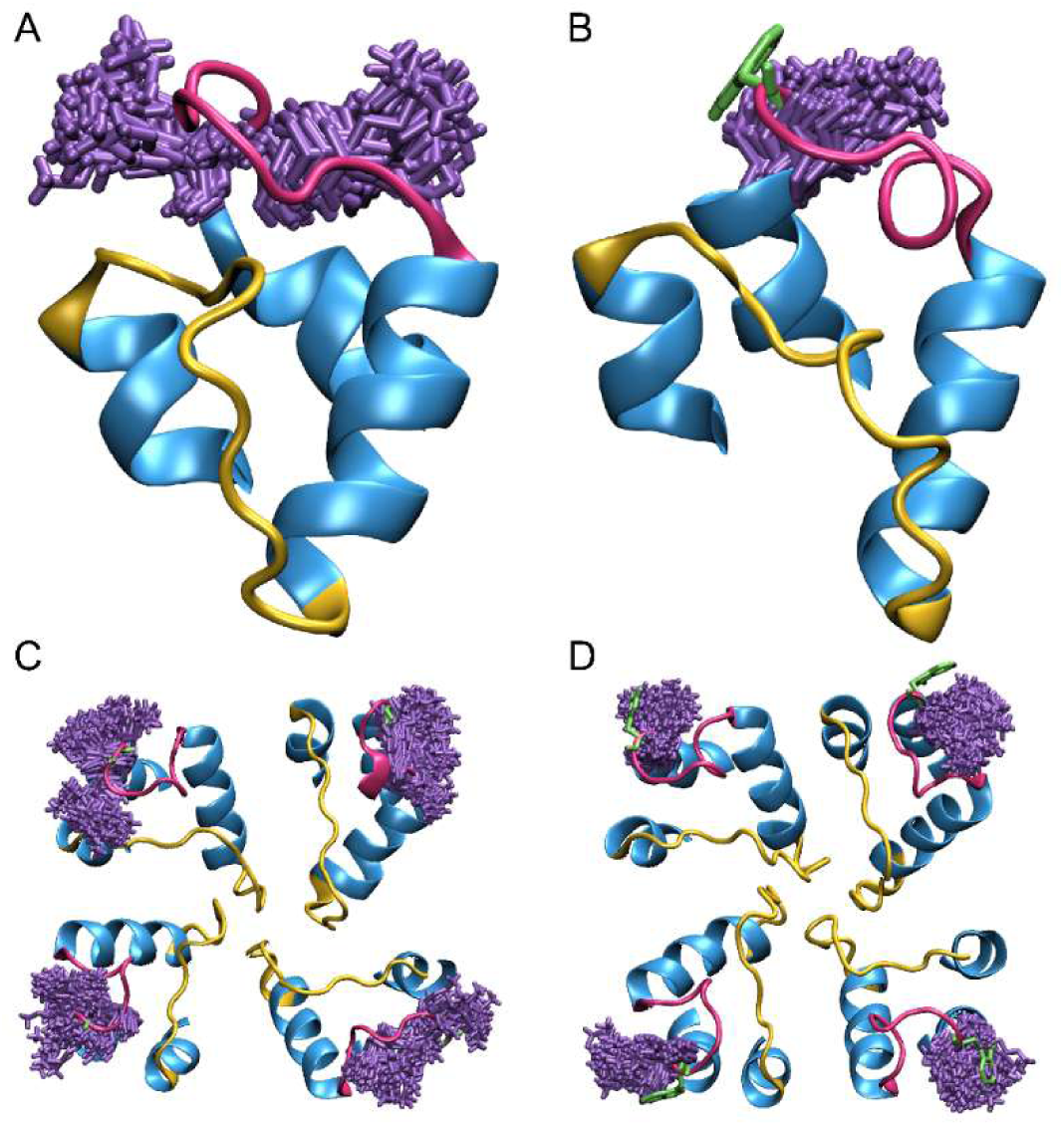
K255 has access to more conformational states in the WT structure. K255 (purple) positions are superimposed every 10 ns for all simulations (1.5 µs) of **A.** WT and **B.** G256W channels. Extracellular view of static representative G256 and W256 pore domains with S5 and S6 helices restricted to the extracellular halves and intervening p-loop are shown in **C** and **D**. K255 locations shown in Figure 5 of all subunits superimposed every 10 ns for all simulations (1.5 µs) of **C.** WT and **D.** G256W. The protein ribbon is shown in blue with a yellow selectivity filter and pink turret. A W256 skeletal structure is shown in green in panel **B** and **D**.

The side chain of N258 separated the side chains of E254 and H260 along the turret initially for both systems. G256 flexibility allowed access to additional conformations for all amino acids in the turret (Figure S3). This flexibility enabled N258 to rotate away from E254 and H260, which caused the side chains E254 and H260 to interact for 15.9% of the WT simulation, as opposed to 0.1% in the less flexible G256W turret.

Key hydrogen bonding frequencies that connect the turret to the selectivity filter bridge such as the interaction between H260 and D266 with K283 near the top of the selectivity filter were lower for G256W (Figure 7A). The interaction between E254 and Q286 was similar between structures, likely because, as part of the S5 helix and thus part of an ordered secondary structure, E245 was less affected by the variant. The backbone RMSF also verified that the WT and G256W E254 did not deviate from each other (Figure S5). E254 did however have a 16.7% higher interaction with N258 in the G256W which suggested that the G256W turret may have placed the N258 side chain closer to and increased the strength of interacting with E254 (Figure 7A). The hydrogen bonding network with and including N258 seemed to move the turret slightly near the SF bridge in the G256W structure, closer to Q286 and farther from H260 (Figure 7A). This was indicated by a near 10% increase (Figure 7A) in interaction frequency with Q286 and a 6.9% decrease with H260 for the G256W structure. Denoted by the higher frequencies near W256, the turret adopted a conformation that shifted towards W256 when substituted. Subsequent pulling on the turret from Trp membrane burial indicated by SASA analysis and the distance to membrane center plots provided an explanation for more rigid G256W side chain interactions and the potential shift in conformation (Figure 5).

The hydrogen bonding interaction frequencies between K283 with H260 and D266 were lower for the G256W than in the WT. The backbone RMSF of K283 for the WT (1.0 Å) and G256W (1.1 Å) were similar, which indicated that the turret location and side chain fluctuation were the contributing factors to the change in interaction frequencies. K283’s location at the top of the selectivity filter and on the opposite end of the mobile turret from the variant meant that the shift towards the tryptophan due to its interaction with the membrane may have pulled the turret away from K283. Movie S1 compared the K283 side chain conformations. K283 spent less time interacting with H260 and D266, which was qualitatively associated with more fluctuation within the residues of the canonical selectivity filter. This altered interaction may have contributed to the altered pore dynamics and profile for the G256W structure.

### 2.4 G256W selectivity filter has increased fluctuation

Compared to the WT, the G256W selectivity filter had a larger deviation near the intracellular mouth of the selectivity filter. Average RMSD between opposing subunit selectivity filter backbone and carbonyl atoms was calculated. Results indicated that G256W simulations have a greater RMSD with at least a medium effect size for four out of the five selectivity filter residues: T277, I278, G279, Y280, but not G281 (Table S3). Figure 9 showed a pictorial understanding of these deviations with 1 ns superimpositions of the canonical selectivity filter carbonyls, which line the pore.^40^ Interestingly, the G256W selectivity filter superimpositions qualitatively exhibited a loss of symmetry around the pore indicative of potentially altered selectivity filter dynamics (Figure 9B).

**Figure 9.**
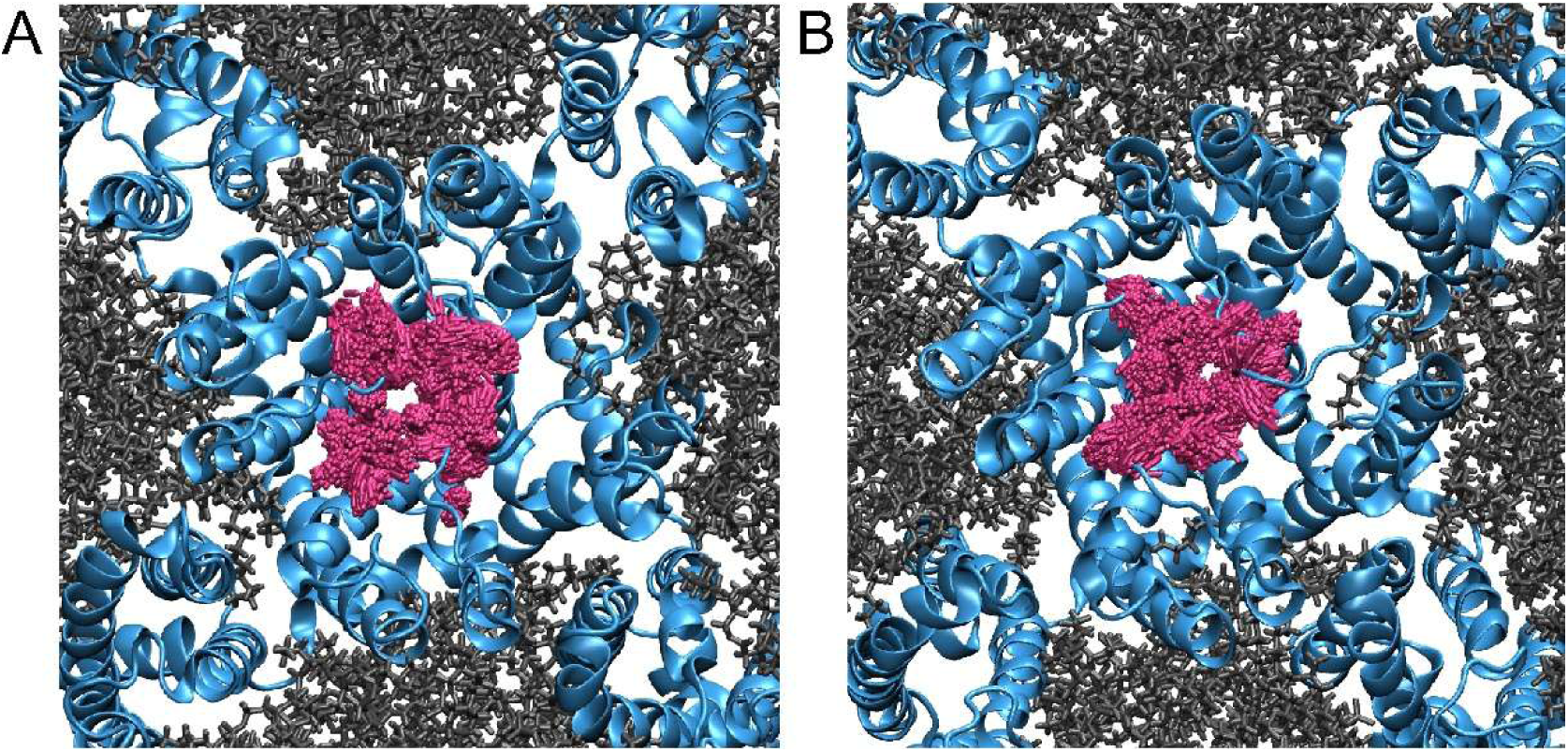
Representative images illustrate altered pore profile and asymmetry that occur with G256W. A. WT and **B.** G256W representative structures of the transmembrane domains (blue). Extracellular view of pore radii using the canonical selectivity filter sequence carbonyls (pink) embedded in a membrane (dark grey) superimposed across all simulations (1.5 µs) for every 1 ns. Intracellular N and C terminal regions are omitted for clarity.

Movie S1 shows representative examples of turret and selectivity filter dynamics for the WT and G256W structures. The movie illustrates more fluctuation for the G256W selectivity filter described quantitatively by an increase in RMSD standard deviation (0.40 Å higher standard deviation for G256W compared to WT) (Figure S3). K283 side chain conformations were farther away from the turret for G256W compared to the WT simulation, consistent with the lower hydrogen bonding frequency measured across the datasets. The critical location of K283 between the turret and selectivity filter makes it a good candidate mediator for effects of G256W substitution on the selectivity filter. This residue is conserved across all KCNQ channels and in KCNQ2. In other inward rectifiers, K283 or the analogous position has been shown to change the selectivity for different ions.^41^ Thus, G256W may impact the selectivity filter dynamics through its disruption of K283 interactions.

Differences in the pore profile were expected to have effects on ion conduction. Ions traverse the pore by coordinating with the selectivity filter carbonyl oxygens that replace the water shell lost and associated energy cost of dehydration.^40,42^ Thus, changes in pore geometry would be expected to impact these exquisitely conserved interactions and be poorly tolerated. The altered size and dynamics exhibited by the G256W pore in our simulations could contribute to the loss of potassium conduction and neuronal hyperexcitability seen when G256W was expressed heterologously in cell lines and in transgenic mice.^5,9^

### 2.5 Effects of the WT:G256W subunit stoichiometry within tetramers on dynamics highlight the turret’s importance for channel function

We conducted additional smaller scale simulations (200 ns) including the full range of potential KCNQ2 subunit stoichiometries predicted to arise in vivo in individuals with heterozygous G256W variants, namely tetramers composed of 3WT:1G256W, 2WT:2G256W (adjacent), 2WT:2G256W (opposing), and 1WT:3G256W subunits. The average ± SD for the backbone RMSD equilibration runs for WT subunit-only and G256W subunit-only tetramers were 4.65 ± 0.92 Å, and 4.11 ± 0.84 Å, respectively. Channels formed by combinations of WT and G256W subunits exhibited similar backbone RMSDs (Figure S6). The RMSD indicates the heteromeric simulations have dynamics between WT and G256W and are proportional to the number of G256W subunits per tetramer, with medium to large effect sizes compared to both (Figure S6; Table S4). Backbone RMSD of the turret region reveals that dynamics of the turret region correspond to the channel stoichiometry and follows the same trend of turret mobility identified earlier: more WT subunits increase turret dynamics (Figure S7A). The selectivity filter dynamics do not follow this same trend to the same degree (Figure S7B). However, Hedges’ g calculations comparing the selectivity filter of the heteromeric simulations to the WT and G256W simulations showed that any tetramer with at least one G256W subunit has SF dynamics more similar to the G256W homotetramer simulation than the WT (Table S5). This is consistent with in vitro electrophysiological recordings showing that G256W variants have dominant-negative effects on conduction.^5^

### 2.6 Additional simulations of turret and selectivity filter bridge pathogenic variants N258Y, H260P, and K283M

To further assess the turret mobility and residue interaction results obtained for G256W, we performed additional MD simulations using three additional variants: N258Y, H260P, and K283M. N258 and H260 were chosen because they represented key amino acids that were part of the turret and selectivity filter bridge connection seen in prior cryoEM-based modeling of turret hydrogen bonding and in the MD analysis (Figure 6; Figure 7). Similarly, interactions with K283 were hypothesized to play a role in turret-selectivity filter dynamics. The electrophysiological consequences of each of these variants have been studied previously; all caused reduced conduction.^41,43^ Missense variants at these positions, although not these specific substitutions, have been reported in individuals with DEE and SLFNE and were not found in the control population, GnomAD.^44–50^ Backbone RMSD for tetrameric simulations of each of these putative pathogenic variants were more similar to G256W than WT (Figure S8; Table S6). Similarly, all variant tetramer selectivity filter dynamics were closer in effect size to G256W than WT, although large effect sizes for both illustrated the different substitutions effects (Table S7).

In the N258Y simulation, even though tyrosine can hydrogen bond, interactions between residue 258 and E254 (81.3%), H260 (25.5%), and Q286 (15.9%) were markedly reduced to less than 15% (Figure 7B). Compared to N258, Y258 appeared more solvent exposed (Movie S2) and was more dynamic (heightened peak in RMSF; Figure S9A). Despite the increased dynamics of the turret, there appeared to be no increased dynamics of the selectivity filter (Figure S10A-B; Movie S2). The loss of 258 interactions may have been compensated for by increased surrounding turret interactions, including H260 to K283, which increased from 41.4% (WT) to 80.9% (N258Y), and H260 to Y284, which increased from <15% (WT) to 45.3% (N258Y; Figure 7A-B). However, these altered interactions resulted in selectivity filter dynamics more like the G256W KCNQ2 simulations than the WT (Table S6). Experimental studies that describe electrophysiology of N258Y indicated current densities of approximately 35% of WT.^43^ Interestingly, previous studies of N258 also revealed the variant N258S that contributes to SLFNE.^51,52^ It is interesting to consider that a S258 substitution, which is closer in structure to asparagine, might be less disruptive of N258 hydrogen bonding interactions than N258Y, thus leading to in the milder SLFNE phenotypic outcome.

P260 movement seemed to cause the turret backbone to change conformations relative to the WT H260 (Movie S3). The change in structure associated with an increase in RMSD between 50 and 100 ns correlated to the movement (Figure S10C-D). However, because proline’s sterics are restrictive, the average RMSD of the turret region was lower than the other two variant simulations. The interaction between the sidechain of H260 and N258 was lost with this substitution, however the interaction with the backbone carbonyl of P260 and the side chain of K283 increased slightly compared to WT (41.4% to 50.5%, Figure 7A and C). N258 also contributed to an interaction connecting to the SF bridge with T287 (increase from <15% in the WT to 22.2% in H260P) but decreased in interacting with Q286 (decrease from 60.6% in the WT to 42.8% in H260P). The RMSF indicated that N258 had increased side chain dynamics, which corresponded to the new interactions (Figure S9). Experimentally, H260P led to current densities of approximately 31% of WT.^43^ The lowered current density was similar in value to the N258Y results, which was consistent with the similar average RMSDs observed for the selectivity filter (N258Y: 1.53 Å, H260P: 1.55 Å, Figure S10).

Electrophysiological studies found that the K283M variant changed the KCNQ2 relative cation selectivity profile.^41^ In simulations of K283M, N258 showed increased hydrogen bonding frequencies with E254 and with T287 (Figure 7A and D). This was consistent with the overall lowered dynamics in the region of N258 compared to the N258Y and H260P variant simulations (Figure S9; Movie S4). However, a slight continuing increase in RMSD for the turret over the time studied indicated that more dynamic changes might be captured by lengthier simulation times (Figure S10 E-F).

Overall, the briefer simulations of varying KCNQ2 WT and G256W stoichiometric tetramers and additional turret substitutions complement the view emerging from the longer simulations comparing WT and G256W homotetramers. The simulations of coassembled WT and G256W subunits are relevant to naturally occurring heterozygous variants and suggested that as few as one G256W subunit per tetramer can impact the selectivity filter and can create similar structural effects as the G256W homotetramer. The additional turret missense variant simulations revealed that single site substitutions can disrupt WT bonding interactions throughout the turret and with the selectivity filter bridge. Recent studies showed that heterozygous G256W variants suppress KCNQ2 channel current density to 41% of WT in heterologous cells with conditions analogous to the experimental data for N258Y and H260P mentioned earlier.^5,43^ G256W’s similar reduction in current density compared to the additional turret variant simulations might be explained by all substitutions’ similar mechanisms of altering key turret-selectivity filter bridge interactions seen in the additional turret variant simulations. In all three simulations, interactions with K283 were impacted, which is crucially situated to effect changes in the selectivity filter. N258 and H260P impact conduction slightly more than G256W, which might be due to their positions in the network closer to the selectivity filter.

## 3. CONCLUSION

Studies of the biophysical consequences of pathogenic KCNQ2 variants may yield new insights into channel mechanisms and potential therapeutic avenues. Previously, Abreo et al. combined analysis of cryoEM structural models and functional studies of the pathogenic variant, G256W. Abreo et al. found that G256W caused loss-of-function by reducing conductance without changing voltage-dependence, a pattern consistent with dominant blocking of the ion pore. The authors suggested that a hydrogen bonding network linking the turret, including G256, might help stabilize the selectivity filter in an open state. We have further tested that idea through all-atom MD simulations. We reported similar backbone deviation in the turret region for WT and G256W, but reduced side chain flexibility for the G256W turret. We identified W256 sidechain migration toward the lipid membrane and steric and electrostatic restriction of K255 mobility as prominent potential contributors to reduced G256W turret flexibility. Observed hydrogen bonding frequencies that involved turret residues revealed interactions that overlapped with but also differed from those predicted from Abreo et al.’s analysis of cryoEM data. The prior interpretation was that a network of hydrogen bonds in the WT turret might be essential for stabilizing the pore, acting via the selectivity filter bridge segment like an extracellular “flying buttress”. Thus, it was unexpected that G256W did not appear to destabilize the pore by destabilizing the turret; instead, G256W increased turret stability in an abnormal conformation due to novel interactions (including: W256 lipid interaction, K255 restricted mobility). These abnormal interactions disfavor ion conduction.

It is very challenging to model a protein as large and complex as a KCNQ2 tetramer at the very high spatial and temporal resolution important for ion conduction. Our study was performed in the absence of an imposed membrane potential, and in a simplified all 1-palmitoyl-2-oleoylphosphatidylcholine (POPC) membrane lipid environment. Inner leaflet phosphatidyl inositol-4,5-bisphosphate (PIP2), which is required for channel opening in vivo, was omitted.

The x and y dimensions (∼15 nm) of our membrane unit cell were similar to the diameter of nanodiscs used in recent cryoEM studies of PIP2 binding sites near the ion pore. The HA-HB linker, which we omitted and which has been unresolved by cryoEM studies to date, interacts with membrane PIP2 to control channel gating, though possibly at a distance from the pore greater than the radius of the nanodiscs and MD unit cell.^53,54^ It will be interesting in future simulations to model larger membrane areas, include the HA-HB linker, and attempt to study PIP2 interactions of the entire channel.

Human pathogenic variants are heterozygous, and, owing to random association, cells are thought to express channels contain a mix of WT and variant subunits. Our modeling of KCNQ2 channels containing 1 to 4 G256W subunits per channel showed similar impacts on pore dynamics, indicative of the variant’s known dominant negative effect on current density. In sum, this study provides additional support for the idea that a bonding network between the turret and selectivity filter helps enable KCNQ2 pore conduction and highlights additional residues that could be investigated by site-directed mutagenesis and electrophysiological analysis.

## 4. EXPERIMENTAL PROCEDURES

### 4.1 System Preparation

We obtained the KCNQ2-CaM structure in the apo state (PDB: 7CR3).^14^ The resolved structural model lacked several portions of the intact KCNQ2 polypeptide: the distal intracellular N-terminus (residues 1-70) and C-terminus (596-872), the extracellular S3-S4 loop (184-193), the HA-HB intracellular linker (367-535). Using UCSF Chimera MODELLER, the S3-4 linker was modeled (Figure 1A). For simplicity and due to the uncertainty in modeling in regions with so many amino acids, the intracellular termini and HA-HB loop were not added. The distal HA helix and proximal HB helix ends were terminated, and not *in silico* connected. G256W substitutions were introduced manually to all subunits by modifying the PDB structure and visualized to confirm no overlap with other atoms. The tryptophan was built according to the AMBER standard coordinates provided by the force field, ff14SB. Using the CHARMM-GUI Membrane Builder, a POPC (1-palmitoyl-2-oleoylphosphatidylcholine; 359 phospholipids total) lipid bilayer was generated around the protein.^55–66^ The structure and newly placed residue substitutions were manually checked in VMD to ensure no overlapping atoms. The lipids and protein were solvated above and below the membrane central plane and around the intracellular or extracellular portions of the protein with 0.15 M KCl in two TIP3P water layers with a minimum height of 22.5 Å. The unit cell dimensions were 151 x 151 x 156 Å. An image of the unit cell set up is shown (Figure S1). The total atoms for the WT simulation were 240,840 atoms, omitting changes due to residue substitutions.

### 4.2 Molecular Dynamics Simulations

Triplicate WT and G256W 750 ns simulations were performed using Amber24 with ff14SB and Lipid17 force fields.^67–71^ The initial structure was minimized with 25,000 maximum minimization steps, first with 10,000 steps of steepest descent before switching to conjugate gradient minimization. We restrained the solute with a restraint weight of 5 kcal/mol/Å^2^. We heated from 0 K to 100 K linearly over 5 ps NVT and then 100 K to 310 K NPT over 100 ps. During heating, the SHAKE algorithm was applied with a 2 fs timestep and the solute was restrained with a weight of 10 kcal/mol/Å^2^.^72^ The structure was equilibrated over 5 ns NPT without restraints at 310 K. Production runs were performed in triplicate each for 750 ns for both WT and G256W, although the first 250 ns was excluded from analysis as equilibration based on RMSD. To determine the time at which RMSD values plateaued to reach a stable structure, energy stabilization criteria were calculated using the linear regression of the RMSD plots to ensure the production run slope was near 0 (between -0.0001 and 0.0001). In all simulations, the linear regression of the production run reached these criteria around 250 ns, so for clarity and convenience, the production run started at 250 ns. For all simulations, a nonbonded cutoff of 10 Å and periodic boundary conditions were applied. A total of 1.5 µs for WT and G256W simulations were analyzed.

### 4.3 Analysis

Root mean square deviation (RMSD) analysis was performed on the backbone atoms of KCNQ2 compared to the same initial structure for WT and G256W over the full trajectory. RMSD plots were smoothed using the Bezier interpolation in Gnuplot. Root mean square fluctuation (RMSF) of the average fluctuation per residue using heavy atoms of the backbones as well as the backbones plus side chains were calculated. The solvent accessible surface areas (SASA) of G256 and W256 were measured using CPPTRAJ and reported as a percentage with 95% confidence intervals which were normalized to the total surface areas of glycine and tryptophan.^73,37^

*MDAnalysis* was used to calculate the vertical distance to center membrane plots averaging all simulations over the 500 ns triplicate trajectories and plotted with a running average window of 50 frames (thus, the first and last 50 frames are omitted). The HydrogenBondAnalysis module was used (atom distance cut-off, 3.5 Å; 120° angle cut off) to calculate the frequency of hydrogen bonding interactions.^74–76^

Images and movies were produced using Visual Molecular Dynamics (VMD),^77^ Chimera,^78,79^ ChimeraX,^80–82^ and Mol*.^83^

### 4.4 Statistics

For SASA and the vertical distance to membrane center plots, where calculated data points are present in each frame, a Welch’s t-test was performed.^84,85^ For calculations comparing values across replicate simulations (e.g., RMSD, RMSF), where the sample size was six (triplicate simulations of WT and G256W), Hedges’ g magnitudes were used due to its strength in assessing differences between conditions with small sample sizes.^86^ We interpreted Hedges’ g magnitude effect sizes according to Cohen’s rules: small < 0.2, medium 0.2 < x < 0.5, or large > 0.8.^87^

## DATA AVAILABILITY STATEMENT

All data discussed in the paper can be shared upon request by contacting the corresponding authors: Edward C. Cooper, MD PhD and Bill R. Miller III, PhD.

## SUPPLEMENTAL MATERIAL

Supplemental material is provided in a separate document and includes data and other visualizations not shown in the main text.

## ACKNOWLEDGEMENTS

Our computational resources were provided partly by the MERCURY consortium under NSF grant CHE-1229354 (BRM). This work was partly supported by the NINDS Epilepsy Center without Walls U54 NS108874 (ECC).

Molecular graphics images were produced using the UCSF Chimera package from the Resource for Biocomputing, Visualization, and Informatics at the University of California, San Francisco (supported by NIH P41 RR-01081).

Molecular graphics and analyses performed with UCSF ChimeraX, developed by the Resource for Biocomputing, Visualization, and Informatics at the University of California, San Francisco, with support from National Institutes of Health R01-GM129325 and the Office of Cyber Infrastructure and Computational Biology, National Institute of Allergy and Infectious Diseases.

