## Supplemental Information for "Molecular dynamics of the pathogenic KCNQ2 variant G256W reveal mechanisms of channel dysfunction in epileptic encephalopathy"

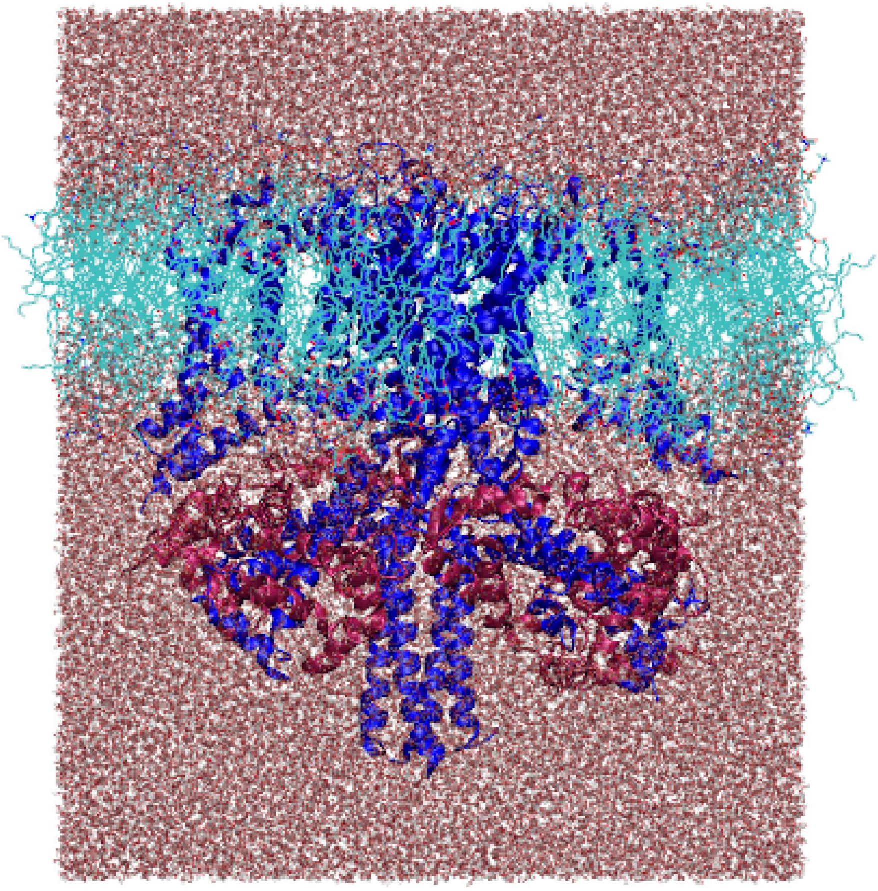

**Figure S1. Simulation unit cell is solvated with TIP3P water and contains the KCNQ2/CaM complex embedded in a lipid bilayer**

Side view of the solvated simulation unit cell, with KCNQ2 homotetramer (blue), Calmodulin (magenta), POPC lipid bilayer (teal), and TIP3P water (red and white).

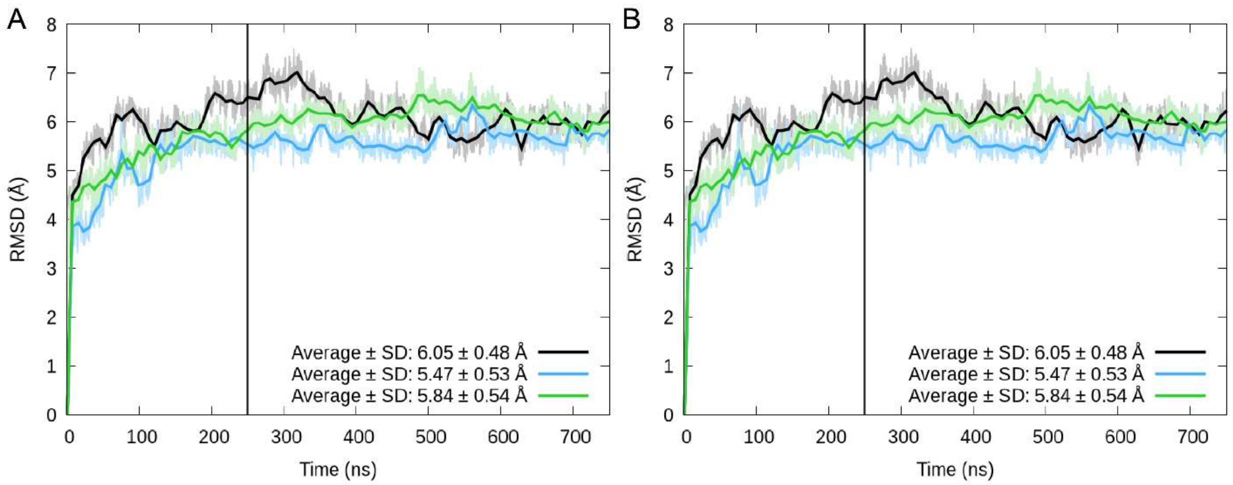


**Figure S2. Root mean square deviation displays structure stabilization after 250 nanoseconds**

Root mean square deviation averaged across all subunits for **A.** WT and **B.** G256W 750 ns simulations. Lines for replicates one (black), two (blue), and three (green) illustrate the full structure RMSD traces (faint) and smoothed average lines (bolded) per simulation. Legend displays the calculated average and standard deviation for each replicate simulation. The first 250 ns was excluded from future analyses as time allowed for the equilibration of the structure. The vertical line indicates the start of the production MD for analysis.

| | WT | | G256W | |
|---|---|---|---|---|
| Simulation time | Average (Å) | Standard deviation (Å) | Average (Å) | Standard deviation (Å) |
| 0 - 250 ns | 5.39 | 0.68 | 5.62 | 0.67 |
| 250 - 750 ns | 5.98 | 0.38 | 6.25 | 0.52 |

**Table S1. Reduced standard deviation after 250 nanoseconds indicates more stability in structure for the 500 nanosecond production runs**

Average and standard deviation of the WT and G256W trajectory RMSD comparing the initial 250 ns from each trajectory with the final 500 ns of each 750 ns trajectory.

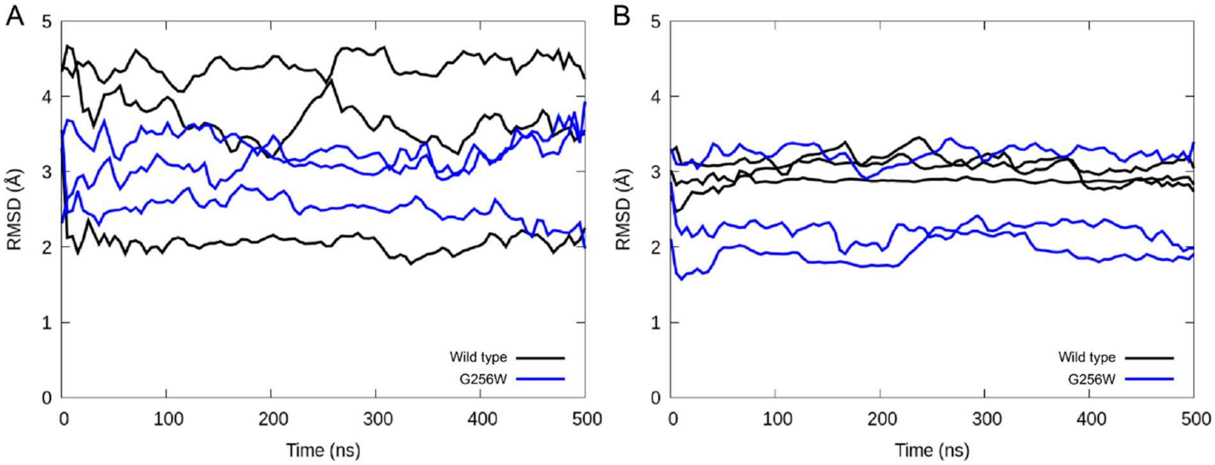


**Figure S3. Wild type KCNQ2 tetramers have more dynamic turret regions and more stable selectivity filters than G256W tetramers**

RMSD results of the **A.** turret and **B.** selectivity filter backbone segment residues for the triplicate WT and G256W simulations. For the turret, WT simulations had an average RMSD ± SD of 3.38 ± 1.02 Å. The G256W simulations average RMSD and SD was 2.99 ± 0.44 Å. For the selectivity filter, the average RMSD and SD was 3.01 ± 0.18 Å (WT) and 2.46 ± 0.59 Å (G256W).

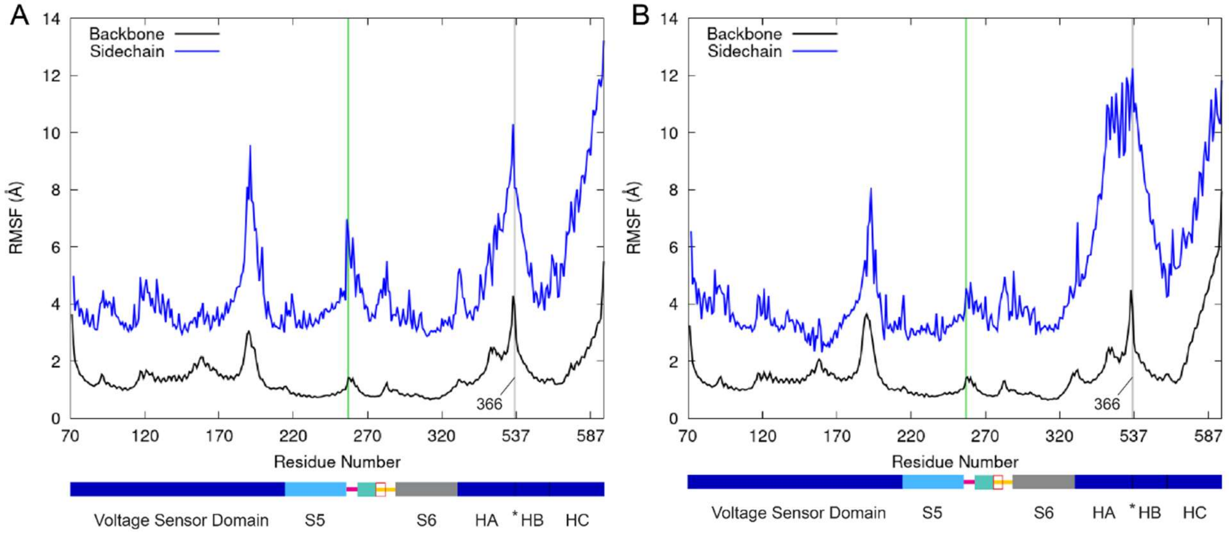


**Figure S4. Inclusion of sidechain atoms highlight a broader range of structural fluctuation**

Average root mean square fluctuation per residue over all subunits comparing calculations including backbone atoms only with backbone and sidechain atoms of **A.** wild type and **B.** G256W. Green lines indicate residue 256, and grey lines mark the gap between residues 366 and 539 that correspond to the unresolved HA-HB loop. Schematic diagrams are shown below the graphs.

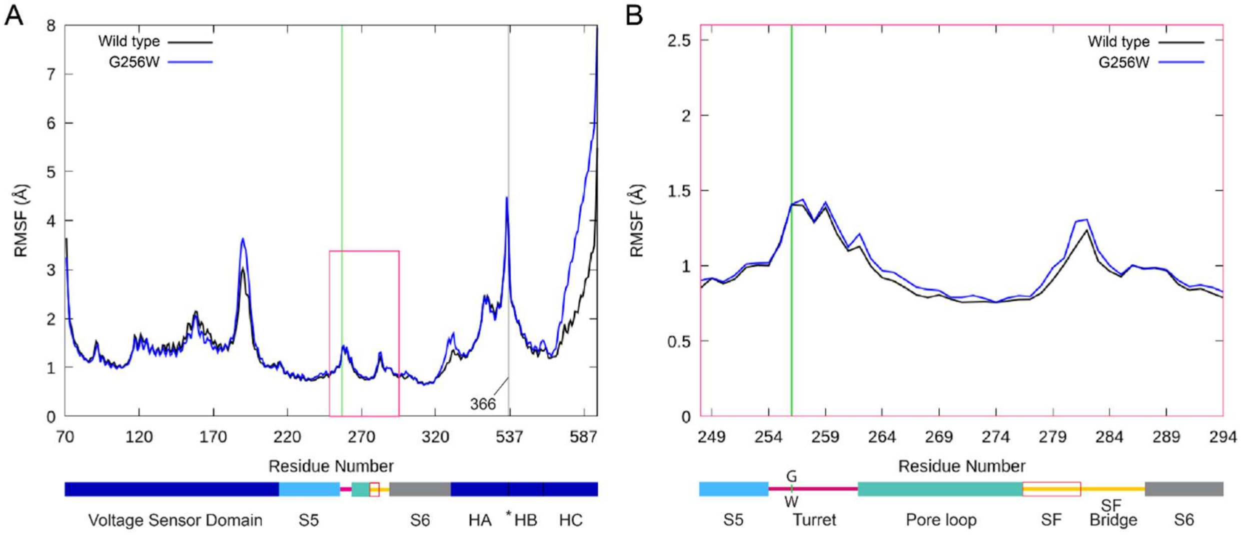


**Figure S5. Backbone root mean square fluctuation is similar for the WT and G256W, with the greatest differences in the S3-S4 loop and HC helix**

Root mean square fluctuation per residue averaged over all subunits for all simulations only including backbone atoms. A schematic representation of the domains of KCNQ2 is shown below the graph with colors corresponding to the domains in Figure 1A. The green line marks 256 in both panels. **A.** Comparison of the wild-type (black) and G256W (blue) structures of KCNQ2 residues. The grey line at residue 366 indicates a jump in sequence to residue 536 corresponding to a poorly resolved HA-HB loop. **B.** A zoom-in on the pore domain region boxed in pink in panel A depicting a less flexible G256W turret region, especially near W256.

| Residue | Backbone Hedges' g magnitude | Sidechain Hedges' g magnitude |
|---|---|---|
| Lys 255 | 0.12 | 0.56 |
| Gly 256 Trp | 0.01 | 1.33 |
| Glu 257 | 0.12 | 0.21 |
| Asn 258 | 0.03 | 0.20 |
| Asp 259 | 0.12 | 0.14 |
| His 260 | 0.24 | 0.31 |
| Phe 261 | 0.18 | 0.38 |
| Asp 262 | 0.53 | 0.42 |

**Table S2. Turret RMSF including side chains is more affected by the G256W substitution than only backbone atoms**

Hedges' g magnitude comparing WT and G256W of each residue in the turret and corresponding atoms showing larger Hedges' g magnitudes for the side chains, suggesting they are more affected by the G256W mutation than the backbone atoms.

https://youtu.be/1jzvTEXjRQ8

**Movie S1. K283 and the selectivity filter fluctuate less in the WT than the G256W structure**

Movie of a representative 500 ns WT and G256W simulation performed on an intact channel tetramer. The extracellular portion of one subunit is shown and colored as in Figure S8. Sidechains omitted except for G256 (left, green, turret), W256 (right, green, turret), and K283 (green, selectivity filter bridge).

| Residue | WT (Å) | G256W (Å) | Hedges' g magnitude |
|---|---|---|---|
| T277 | 0.10 | 0.12 | 0.57 |
| I278 | 0.28 | 0.37 | 0.83 |
| G279 | 0.22 | 0.34 | 0.82 |
| Y280 | 0.23 | 0.32 | 0.62 |
| G281 | 0.34 | 0.32 | 0.10 |

**Table S3. Average RMSD between opposing subunits of selectivity filter indicate greater deviation for the G256W simulations for four out of five residues**

Average root mean squared deviation calculated of the backbone atoms including the carbonyl oxygens between subunits on opposite sides of the pore. The Hedges' g magnitude in the far-right column describes effect sizes between WT and G256W. All residues except for G281 have greater deviation for the G256W simulations.

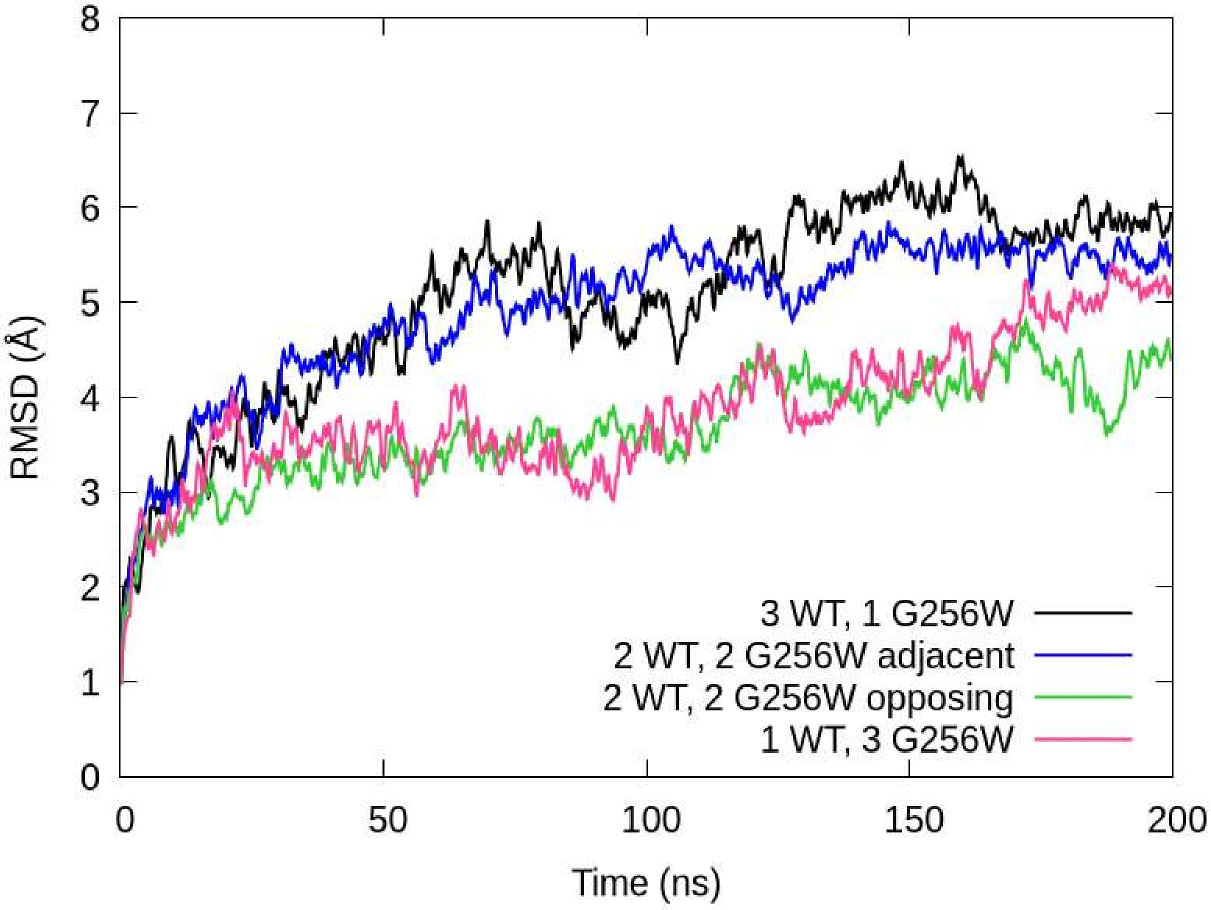


**Figure S6. Backbone RMSD of the WT:G256W stoichiometric simulations are similar to WT and G256W equilibration RMSD**

Backbone RMSD comparison across 200 ns WT:G256W simulations (see legend). Average ± SD RMSD values (listed top to bottom in the legend) are 5.05 ± 1.01 Å, 4.91 ± 0.81 Å, 3.65 ± 0.58 Å, and 3.86 ± 0.73 Å, respectively. These simulations are most comparable to the equilibration of the WT and G256W full simulations (Figure S2).

| Simulation | Hedges’ g magnitudes compared to 4 WT tetramer | Hedges’ g magnitudes compared to 4 G256W tetramer |
|---|---|---|
| 3 WT, 1 G256W | 0.43 | 1.07 |
| 2 WT, 2 G256W (adjacent) | 0.29 | 0.95 |
| 2 WT, 2 G256W (opposing) | 1.16 | 0.58 |
| 1 WT, 3 G256W | 0.90 | 0.32 |

**Table S4. Medium to large effect sizes of RMSD between the WT:G256W stoichiometric simulations and both WT and G256W suggest conformations proportional to the ratio of WT:G256W and between the WT and G256W simulations**

Hedges’ g magnitudes comparing the backbone RMSD between WT or G256W equilibration periods and each additional WT:G256W subunit brief simulation.

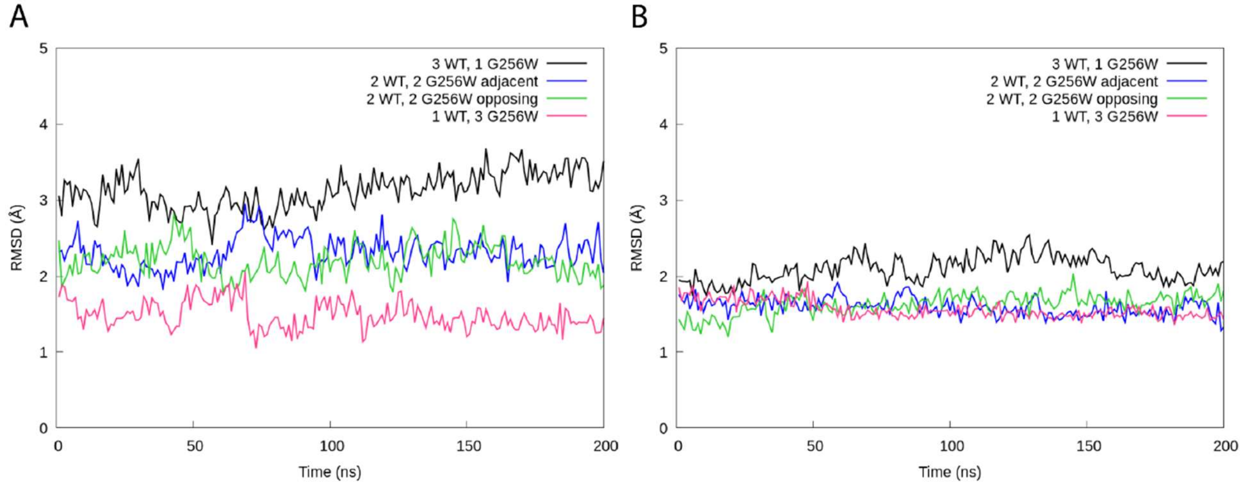


**Figure S7. A range of differences in the turret region among differing WT:G256W stoichiometric simulations cause similar selectivity filter dynamics**

Backbone root mean square deviation of the **A.** turret and **B.** selectivity filter backbone atoms for each WT:G256W simulation (see legend). Average ± SD of turret RMSD (listed top to bottom in the legend): 3.00 ± 0.21 Å, 2.78 ± 0.38 Å, 2.06 ± 0.19 Å, and 1.88 ± 0.27 Å. Average ± SD of selectivity filter RMSD (listed top to bottom in the legend): 2.10 ± 0.13 Å, 1.62 ± SD 0.18 Å, 1.84 ± 0.12 Å, and 1.50 ± 0.07 Å.

| Simulation | Hedges' g magnitudes compared to 4 WT tetramer | Hedges' g magnitudes compared to 4 G256W tetramer |
|---|---|---|
| 3 WT, 1 G256W | 8.7 | 1.7 |
| 2 WT, 2 G256W (adjacent) | 7.6 | 1.5 |
| 2 WT, 2 G256W (opposing) | 6.6 | 1.1 |
| 1 WT, 3 G256W | 5.1 | 0.6 |

**Table S5. The presence of at least G256W subunit causes SF dynamics more similar to G256W than WT**

Hedges' g magnitudes compare the selectivity filter RMSD between WT or G256W and each WT:G256W stoichiometric simulation. G256W has lower Hedges' g magnitudes compared to WT for all brief simulations regardless of WT:G256W ratio.

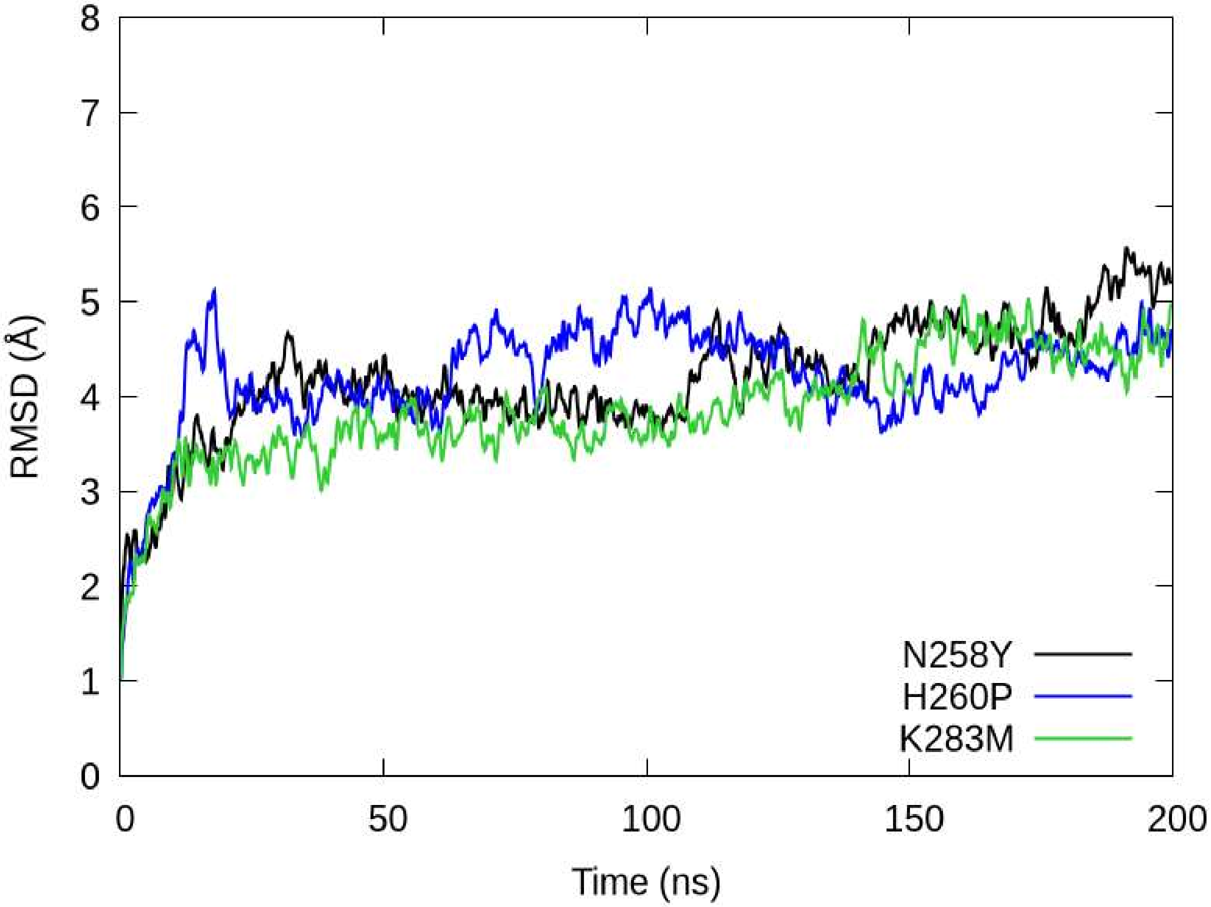


**Figure S8. Backbone RMSD of the additional variant simulations reveal G256W-like dynamics**

RMSD of the full protein structure backbone atoms over time for each simulation (see legend). Average and standard deviation RMSD (listed top to bottom in the legend): 4.19 ± 0.63 Å, 4.22 ± 0.55 Å, 3.87 ± 0.60 Å, respectively.

| Variant | Hedges’ g magnitudes compared to 4 WT tetramer | Hedges’ g magnitudes compared to 4 G256W tetramer |
|---|---|---|
| 4 N258Y tetramer | 0.53 | 0.09 |
| 4 H260P tetramer | 0.50 | 0.13 |
| 4 K283M tetramer | 0.90 | 0.31 |

**Table S6. Variant systems have smaller effect sizes compared to G256W than WT indicating a more pathological conformation**

Hedges’ g magnitudes comparing the backbone RMSD between WT or G256W equilibration periods and each additional variant brief simulation.

| Variant | Hedges’ g magnitudes compared to 4 WT tetramer | Hedges’ g magnitudes compared to 4 G256W tetramer |
|---|---|---|
| 4 N258Y tetramer | 8.3 | 1.7 |
| 4 H260P tetramer | 8.1 | 1.6 |
| 4 K283M tetramer | 9.3 | 2.0 |

**Table S7. Variant brief simulation selectivity filter dynamics are more like G256W**

Hedges’ g magnitudes comparing the selectivity filter RMSD between WT or G256W and each additional variant simulation.

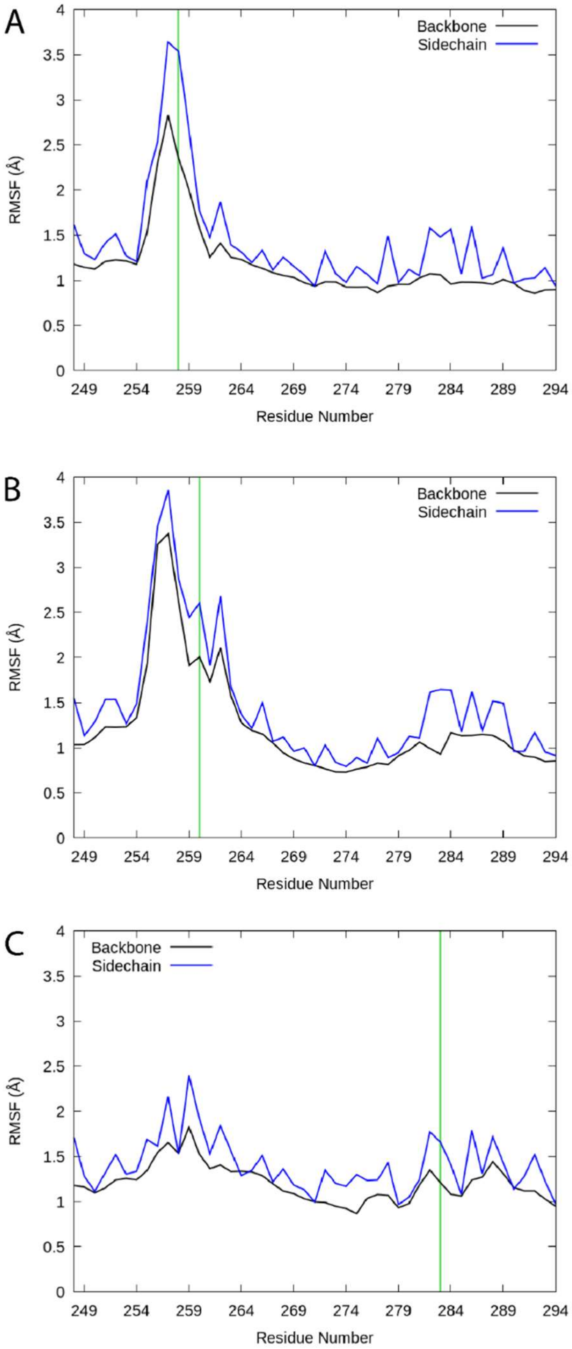


**Figure S9. Key residues in the turret and SF filter alter protein fluctuation**

Per-residue root mean square fluctuation of backbone only (black) and including sidechains (blue) for p-loop residues. RMSF was calculated for **A.** N258Y, **B.** H260P, and **C.** K283M. Green lines indicate the position of substitution.

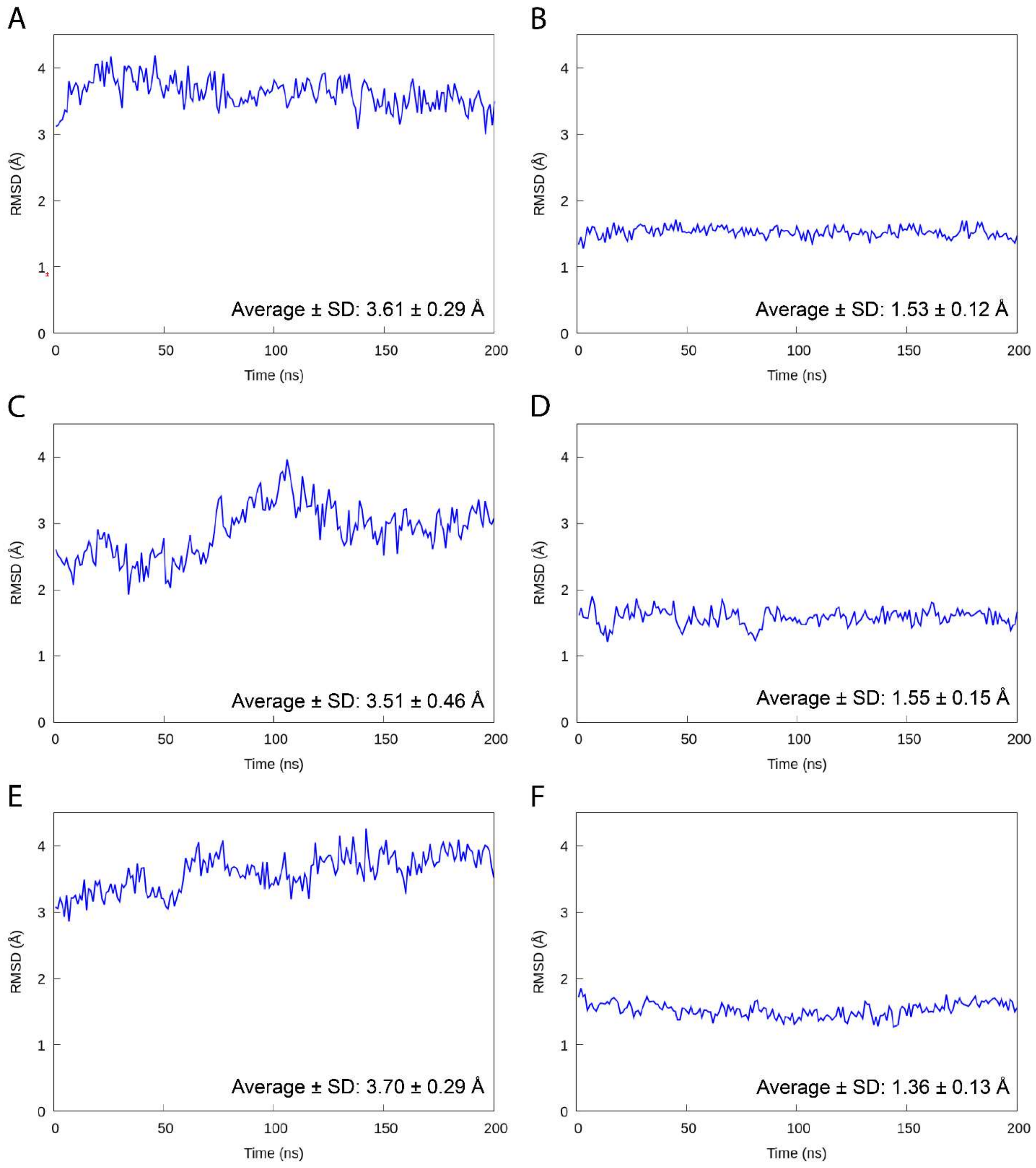


**Figure S10. RMSD of turret and SF for additional simulations indicate how each variant alters dynamics**

Backbone RMSD of the **A.** N258Y turret, **B.** N258Y selectivity filter, **C.** H260P turret, **D.** H260P selectivity filter, **E.** K283M turret, **F.** K283M selectivity filter. Average and standard deviation for each simulation are displayed in its respective plot.

https://youtu.be/XmwcaGO8Xvw

**Movie S2. Dynamic conformational changes in the N258Y simulation**

Movie of 200 ns N285Y simulation performed on an intact channel tetramer. The extracellular portion of one subunit is shown and colored as in Figure S8. Sidechains were omitted except for Y258 (upper left, turret), H260 (upper right, turret), and E254 (lower right, S5) shown. Polar Y258 often rotates toward the solvent.

https://youtu.be/AsumMJnuvL0

**Movie S3. P260 movement away from the SF bridge increases turret mobility**

Movie of 200 ns H260P simulation performed on an intact channel tetramer. The extracellular portion of one subunit is shown and colored as in Figure S8. Sidechains were omitted except for P260 (upper right, turret), Y258 (upper left, turret), and E254 (lower right, S5) shown. P260's flip away from view induces more turret movement.

https://youtu.be/m492FQ-0ZN4

**Movie S4. While overall turret dynamics gradually rise, no single residue displays drastic movement**

Movie of 200 ns K283M simulation performed on an intact channel tetramer. The extracellular portion of one subunit is shown and colored as in Figure S8. Sidechains were omitted except for M283 (upper right, turret), Y258 (upper left, turret), and E254 (lower right, S5) shown. M283 does not interact with H260P or D262 as frequently as K283.
